# Long-read genome assemblies reveals a *cis*-regulatory landscape associated with phenotypic divergence in two sister *Siniperca* fishes

**DOI:** 10.1101/2022.11.09.515789

**Authors:** Guangxian Tu, Qi Chen, Xinshuang Zhang, Ruirun Jiang, Long Zhang, Chengjun Lai, Zhuyue Yan, Yanrong Lv, Shaoping Weng, Li Zhang, Jianguo He, Muhua Wang

**Author notes:** Correspondence: Muhua Wang; Jianguo He.

## Abstract

**Background:** Dissecting the genetic basis of variation in the regulation of gene expression is essential for understanding phenotypic evolution. Structural variants intersecting the *cis*-regulatory elements are found to cause gene expression variation in several developmental genes, resulting in morphological divergence between species. Due to the difficulty of identifying structural variants accurately across the genome, a comprehensive study of impacts of structural variants in *cis*-regulatory divergence of closely related species, especially fish species, is still scarce. Recently identified broad H3K4me3 domains are essential for the regulation of genes involved in several biological processes. However, the role of broad H3K4me3 domains in phenotypic divergence remain poorly understood. *Siniperca chuatsi* and *S. scherzeri* are two closely related fish species diverge in several phenotypic traits, making them an ideal model to study *cis*-regulatory evolution in closely related species.

**Results:** We generated chromosome-level genomes of *S. chuatsi* and *S. scherzeri*. The evolutionary histories of *S. chuatsi* and *S. scherzeri* were studied by inferring the dynamic changes in the ancestral population sizes. The genetic basis of adaptation in *S. chuatsi* and *S. scherzeri* was dissected by performing gene family expansion and contraction analysis and identifying positively selected genes (PSGs). To investigate the role of SVs in *cis*-regulatory divergence of closely related fish species, we identified high-quality SVs between *S. chuatsi* and *S. scherzeri*, as well as H3K27ac and H3K4me3 domains. Integrated analysis revealed that *cis*-regulatory divergence caused by SVs played an essential role in the differentiation of metabolism, skin pigmentation, and immunity between *S. chuatsi* and *S. scherzeri*. Additionally, divergent broad H3K4me3 domains were found to mostly associate with cancer-related genes in *S. chuatsi* and *S. scherzeri* and contribute to their phenotypic divergence.

**Conclusions:** Our analysis reveals SVs play an essential role in *cis*-regulatory variation between the two sister fish species, which in turn contributes to their phenotypic divergence. The divergence of broad H3K4me3 domains contributes to phenotypic divergence between closely related species. Additionally, the association of broad H3K4me3 domains and cancer-related genes has an ancient origin.

## Introduction

Evolutionary biologists have long sought to elucidate which genetic variants contribute to the evolution of morphological diversity [1]. The accurate and robust regulation of gene expression is critical for the development of living organisms and is a common source of evolutionary change [2]. The variation in gene expression regulation is considered to be a common source of evolutionary change [3]. Accumulated empirical evidence has shown that variation in the regulation of gene expression contributes to morphological divergence between species and populations within the same species, which may in turn result in the adaptation and speciation of various species [4, 5]. Thus, dissecting the genetic basis of variation in the regulation of gene expression is essential for understanding phenotypic evolution [3].

Gene expression variation is largely caused by mutations in *cis*-regulatory elements (CREs), which are collections of transcription binding sites and other non-coding DNA required for transcription activation [6, 7]. Several types of mutations that are responsible for *cis*-regulatory divergence between species have been identified. Nucleotide substitutions caused by point mutations are commonly considered to cause divergence in *cis*-regulatory activity [3]. In addition, structural variants (SVs) intersecting *cis*-regulatory elements of several developmental regulatory genes were found to cause morphological divergence between species [8, 9]. Recently, the availability of high-quality reference genomes for several species enabled genome-wide analyses of SV roles in *cis*-regulatory divergence, revealing that gene expression is widely impacted by SVs affecting CREs [10, 11]. However, a comprehensive study of the roles of SVs in *cis*-regulatory divergence between closely related species, especially fish species, is still scarce.

Histones at different CREs, including promoters, enhancers, silencers, and insulators, have specific post-translational modifications [12]. Genome-wide profiling of CREs using chromatin immunoprecipitation followed by high throughput sequencing (ChIP-seq), or the recently developed Cleavage Under Targets and Tagmentation (CUT&Tag) strategy revealed the evolution of these regulatory elements [13, 14]. Histone H3 lysine 27 acetylation (H3K27ac) is an important epigenetic mark associated with active enhancers and promoters [15]. Studies of H3K27ac marks in the genome found that enhancers were variable among species and responsible for *cis*-regulatory divergence [16]. In addition to H3K27ac, typical histone H3 lysine 4 trimethylation (H3K4me3), which is restricted to narrow regions (1~2 kb) at 5’ end of the gene, is one of the most recognized epigenetic marks of active transcription [17]. Recent studies revealed that the breadth of enrichment sites plays a key role in determining the functions of H3K4me3 [18]. Broad H3K4me3 domains are essential for the regulation of genes involved in cell identity specification, embryonic development, and tumor suppression [19–21]. Nevertheless, the role of broad H3K4me3 domains in phenotypic divergence remains poorly understood.

Sinipercidae (Perciforms) is a subfamily of freshwater fish comprising three genera, including *Siniperca*, *Coreoperca*, and *Coreosiniperca*. The sinipercids are endemic to the East Asia (China, Korea, Japan, Vietnam), with most species distributed in China. Mandarin fish (*Siniperca chuatsi*) and leopard mandarin fish (*Siniperca scherzeri*) are two closely related species from the genus *Siniperca* [22]. Unlike other sinipercids that are distributed in the river systems of South China, *S. chuatsi* and *S. scherzeri* are widely dispersed from south to north of China, indicating that they are well-adapted to diverse environments in this region [23]. *Siniperca chuatsi* and *S. scherzeri* diverged in several phenotypic traits. First, the two species are divergent in body length, body width, and skin pigmentation (**Figure 1A and B**) [24]. Second, the growth rate of *S. chuatsi* is substantially higher than that of *S. scherzeri*, while *S. chuatsi* is more susceptible to disease than *S. scherzeri* [25]. Third, although sinipercids are typical innate and obligate piscivores that solely feed on live fry of other fish species, *S*. *scherzeri* easier accept dead prey fish or artificial diets than *S. chuatsi* [26]. The strong phenotypic divergence makes *S. chuatsi* and *S. scherzeri* an ideal model to study *cis*-regulatory divergence in closely related species.

**Figure 1.**
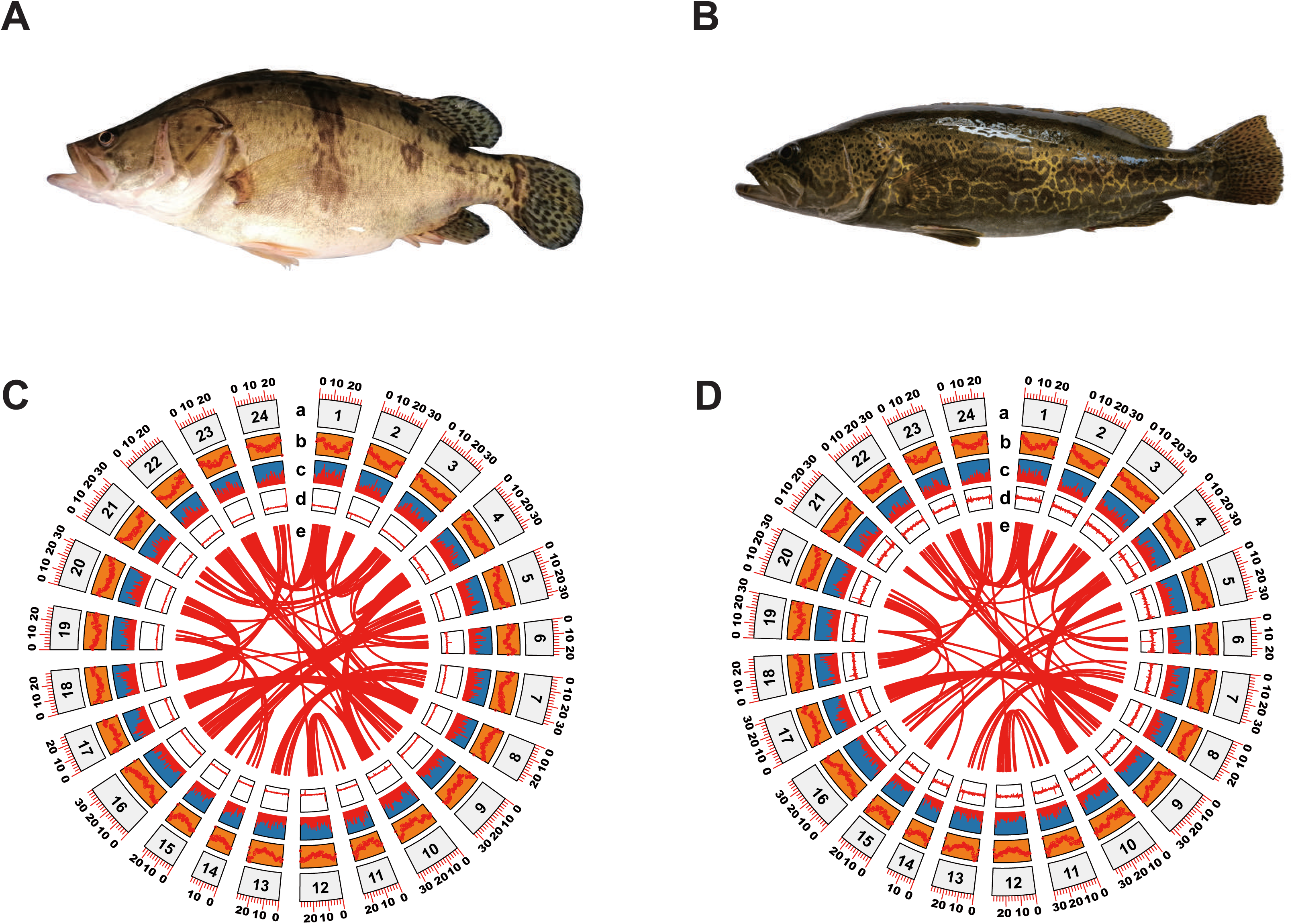
Genome assemblies of *S. chuatsi* and *S. scherzeri*. A large morphological divergence exists between *S. chuatsi* (**A**) and *S. scherzeri* (**B**). Concentric circles show structural, functional, and evolutionary aspects of *S. chuatsi* (**C**) and *S. scherzeri* (**D**) genomes. a. chromosome number; b. repeat density; c. gene density; d. GC content; e. paralogous relationships within the genome.

Here, we used Nanopore sequencing and Hi-C data to assemble the high-quality genomes of *S. chuatsi* and *S. scherzeri*. The evolutionary histories of *S. chuatsi* and *S. scherzeri* were studied by inferring dynamic change of ancestral population sizes. The genetic basis of adaptation in *S. chuatsi* and *S. scherzeri* was dissected by performing gene family expansion and contraction analysis and identifying positively selected genes (PSGs). To investigate the role of SVs in *cis*-regulatory divergence of closely related fish species, we identified high-quality SVs between *S. chuatsi* and *S. scherzeri*, as well as H3K27ac and H3K4me3 domains. Integrated analysis revealed that gene expression variation caused by SVs affecting *cis*-regulatory regions plays an essential role in phenotypic divergence between *S. chuatsi* and *S. scherzeri*. Broad H3K4me3 domains contribute to phenotypic divergence between *S. chuatsi* and *S. scherzeri*. Additionally, divergent broad H3K4me3 domains were mostly associated with cancer-related genes in *S. chuatsi* and *S. scherzeri*, suggesting that the association between this special domain and cancer-related genes has an ancient origin.

## Results

### Chromosome-level genome assembly of *S. chuatsi* and *S. scherzeri*

The genomes of *S. chuatsi* and *S. scherzeri* were sequenced using a combination of Nanopore and Illumina shotgun sequencing. A total of 100.7 Gb of Nanopore reads and 61 Gb of Illumina reads were obtained for *S. chuatsi*, and 94.3 Gb of Nanopore reads and 46 Gb of Illumina reads were obtained for *S. scherzeri* (**Supplementary Tables 1 and 2**). Based on the *k*-mer distribution of Illumina reads, the genome sizes of *S. chuatsi* and *S. scherzeri* were estimated to be 695.9 Mb and 708.4 Mb, respectively (**Supplementary Fig. 1**). The genomes of *S. chuatsi* and *S. scherzeri* were first assembled into contigs with Nanopore reads using the WTDBG assembler [27]. The contigs were then subjected to error correction with Illumina reads. Contigs of *S. chuatsi* and *S. scherzeri* were scaffolded using proximity ligation data from the respective Hi-C libraries to yield genome assemblies (**Supplementary Figs. 2 and 3; Supplementary Tables 3-5**). The final genome assembly of *S. chuatsi* was composed of 191 scaffolds (contig N50: 21.55 Mb, scaffold N50: 29.96 Mb) assembled into 24 pseudomolecules, resulting in a total assembly size of 716.35 Mb. The final genome assembly of *S. scherzeri* consisted of 252 scaffolds (contig N50: 16.04 Mb, scaffold N50: 30.49 Mb) assembled into 24 pseudomolecules with a total assembly size of 740.54 Mb (**Fig. 1C and D; Table 1**). Benchmarking Universal Single-Copy Orthologs (BUSCO) analysis indicated that 97.9% and 98.7% conserved single-copy ray-fin fish (Actinopterygii) genes (odb10) were captured in the *S. chuatsi* and *S. scherzeri* genomes, respectively (**Supplementary Table 6**).

**Table 1.** Genome assembly statistics of S. chuatsi and S. scherzeri.

|  | *S. chuatsi*<br>(This study) | *S. chuatsi*<br>(sinChu7)<br>[28] | *S. scherzeri*<br>(This study) | *S. scherzeri*<br>(sinSch6b)<br>[28] |
| --- | --- | --- | --- | --- |
| Total length<br>(Mb) | 716.34 | 755.06 | 740.54 | 736.22 |
| Number of<br>scaffolds | 191 | 1,156 | 252 | 2,826 |
| Scaffold N50<br>(bp) | 29,956,575 | 30,508,166 | 30,486,584 | 30,166,107 |
| Scaffold N90<br>(bp) | 24,860,938 | 23,880,225 | 27,297,733 | 23,838,788 |
| Number of<br>scaffolds (><br>N90 length) | 21 | 22 | 21 | 22 |
| Number of<br>contigs | 328 | 1,464 | 334 | 23,070 |
| Contig N50<br>(bp) | 21,551,097 | 12,191,788 | 16,036,871 | 83,589 |
| Contig N90<br>(bp) | 3,004,397 | 1,214,209 | 3,383,819 | 16,890 |
| Number of<br>contigs (> N90<br>length) | 42 | 89 | 53 | 9,466 |

We compared the sequence consistency and integrity of our assemblies with previously published genome assemblies of *S. chuatsi* (sinChu7, GCA_011952085.1) and *S. scherzeri* (sinSch6b, GCA_011952095.1) [28]. First, the number of contigs and scaffolds was greatly reduced, and contig N50 was substantially increased in our genome assemblies of *S. chuatsi* and *S. scherzeri* compared to the previously published assemblies (**Table 1**), suggesting the better contiguity of our assemblies. Second, RNA sequencing (RNA-seq) reads of different tissues were aligned to our and previously published genome assemblies. The mapping rates of RNA-seq reads of 9 tissues to our assemblies of *S. chuatsi* and *S. scherzeri* were 92.24% and 96.29%, while the mapping rates of RNA-seq reads to previously published assemblies were 83.37% and 94.28%, respectively (**Supplementary Tables 7 and 8**). Third, Merqury evaluation indicated that the consensus quality values (QVs) of our *S. chuatsi* and *S. scherzeri* assemblies were 36.91 and 35.73, repectively, compared to 32.37 and 30.96 of previously published assemblies, suggesting our assemblies are of high quality (**Supplementary Table 9**) [29]. The genome of *S. chuatsi* carried 196.35 Mb (27.41%) of repeat sequences, and the genome of *S. scherzeri* contained 209.40 Mb (28.28%) of repetitive sequences (**Supplementary Tables 10 and 11**). DNA transposons were the largest class of annotated transposable elements (TEs) in the genomes of the two *Siniperca* species, making up 7.51% and 7.70% of the *S. chuatsi* and *S. scherzeri* genomes, respectively. Protein-coding genes in the genomes of *S. chuatsi* and *S. scherzeri* were identified through a combination of *ab initio*, homology-based, and RNA sequencing (RNA-seq)-based prediction approaches. In total, 29,278 and 29,543 protein-coding genes were identified in the *S. chuatsi* and *S. scherzeri* genomes, respectively (**Supplementary Tables 12 and 13**). BUSCO analysis demonstrated 3,353 (92.1%) and 3,337 (91.7%) complete conserved single-copy ray-fin fish (Actinopterygii) genes (odb10) in the predicted gene models of *S. chuatsi* and *S. scherzeri*, respectively (**Supplementary Fig. 4 and Supplementary Table 14**). A total of 26,623 (90.93%) gene models in the *S. chuatsi* genome and 27,024 (91.47%) gene models in the *S. scherzeri* genome were annotated in at least one of the searched databases (InterPro, eggNOG, KEGG, Swiss-Prot and TrEMBL) (**Supplementary Table 15**).

### Demographic history of *S. chuatsi* and *S. scherzeri*

To investigate the evolutionary history of *S. chuatsi* and *S. scherzeri*, we inferred the histories of ancestral population sizes of these two species using the pairwise sequential Markovian coalescent (PSMC) program [30] (**Fig. 2A**). The ancestral population size of *S. scherzeri* was relatively stable. In contrast, *S. chuatsi* populations expanded in the early Pleistocene (~2 million years ago, Ma) and declined in the early phase of the Mid-Pleistocene Transition (~0.9 Ma), where the duration of the Pleistocene glacial cycles increased from 41 to 100 thousand years ago (ka) [31]. This result suggests that *S. chuatsi* started to colonize new habitats after the prolongation of glacial cycles. The dynamic changes in the ancestral population sizes of *S. chuatsi* and *S. scherzeri* were also inferred using the SMC++ program with genetic variants of 6 wild *S. chuatsi* individuals, 6 wild *S. scherzeri* individuals from south China (Guangdong Province), and 6 wild *S. scherzeri* individuals from north China (Jilin Province) (**Supplementary Table 16**) [32]. SMC++ analysis showed that *S. chuatsi* experienced a population bottleneck at the beginning of the last glacial period (~90 ka), suggesting that the warm temperature in the Eemian interglacial period (129~116 ka) might have facilitated the diversification of this species [33]. *Siniperca scherzeri* from South China have experienced a decline in effective population size at ~400 ka, after the Mid-Brunhes Event (MBE, ~430 ka), where several species experienced dramatic changes in ancestral population sizes (**Figure 2B**) [34–36]. The ancestral population size of *S. scherzeri* from northern China declined at ~300 ka, approximately 100 ka after the population bottleneck of their counterparts from southern China (**Supplementary Fig. 5**).

**Figure 2.**
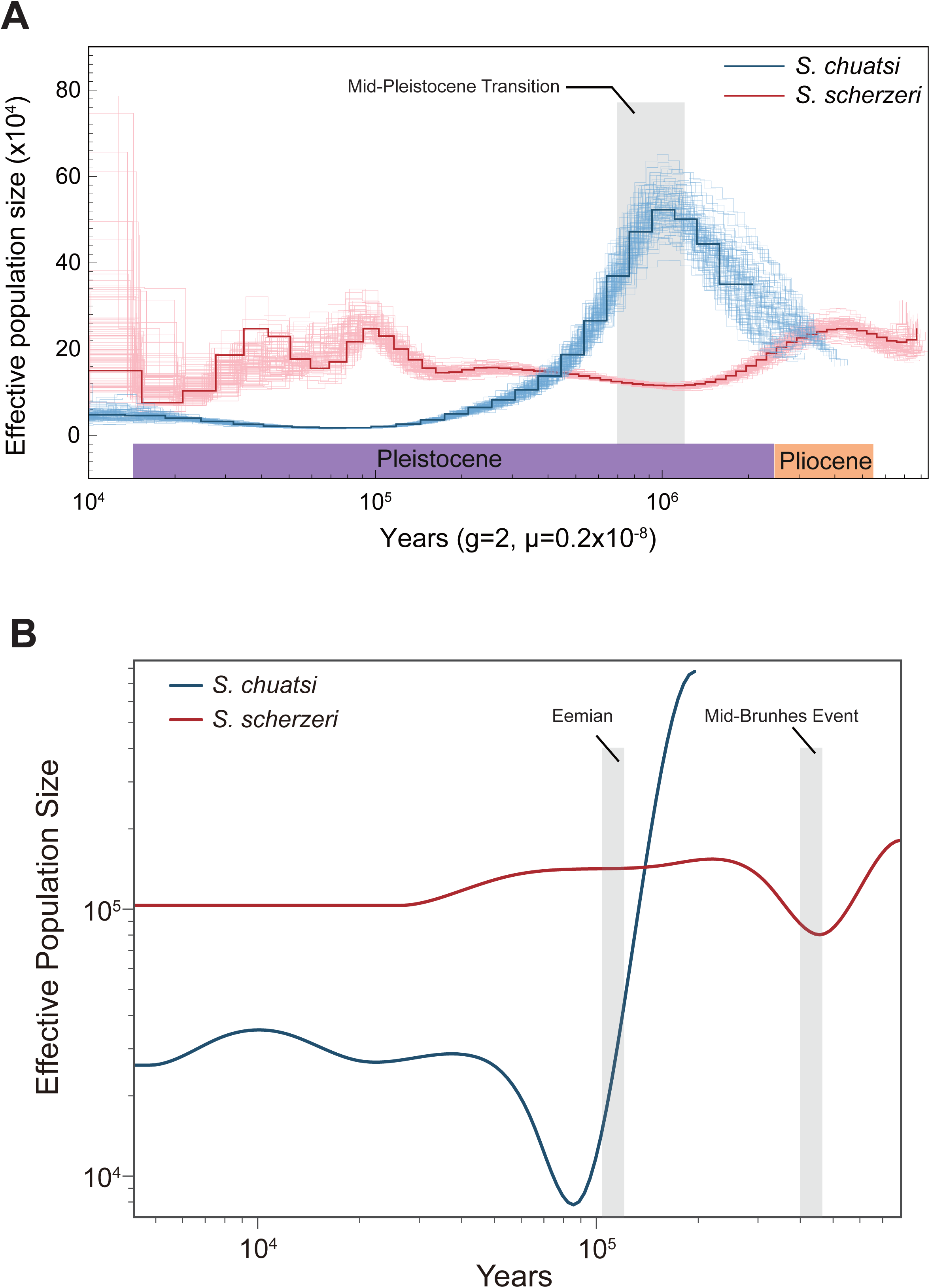
Demographic history of *S. chuatsi* and *S. scherzeri*. **A** Histories of ancestral population sizes of *S. chuatsi* (blue) and *S. scherzeri* (red) inferred using PSMC. **B** Dynamic changes of ancestral population sizes of *S. chuatsi* (blue) and *S. scherzeri* from South China (red) using SMC++.

### Genetic basis of adaptation in *S. chuatsi* and *S. scherzeri*

Gene-family expansion and contraction analysis was performed to dissect the genetic basis of adaptation in *S. chuatsi* and *S. scherzeri*. First, a phylogenomic tree of ten perciform fishes was reconstructed with *Danio rerio* as outgroup (**Figure 3A**). Divergence times were determined among ten perciform fishes. The divergence time of *S. chuatsi* and *S. scherzeri* was estimated to be approximately 14.2 Ma (CI: 1.26-54.45 Ma), corroborating with the results of previous studies [22, 28]. Second, gene-family analysis was performed based on the phylogenomic tree (**Figure 3A**). Compared with other perciform fishes, 72 gene families were expanded, and 384 gene families were contracted in the clade of *Siniperca* (*P* < 0.05) (**Supplementary Data 1**).

**Figure 3.**
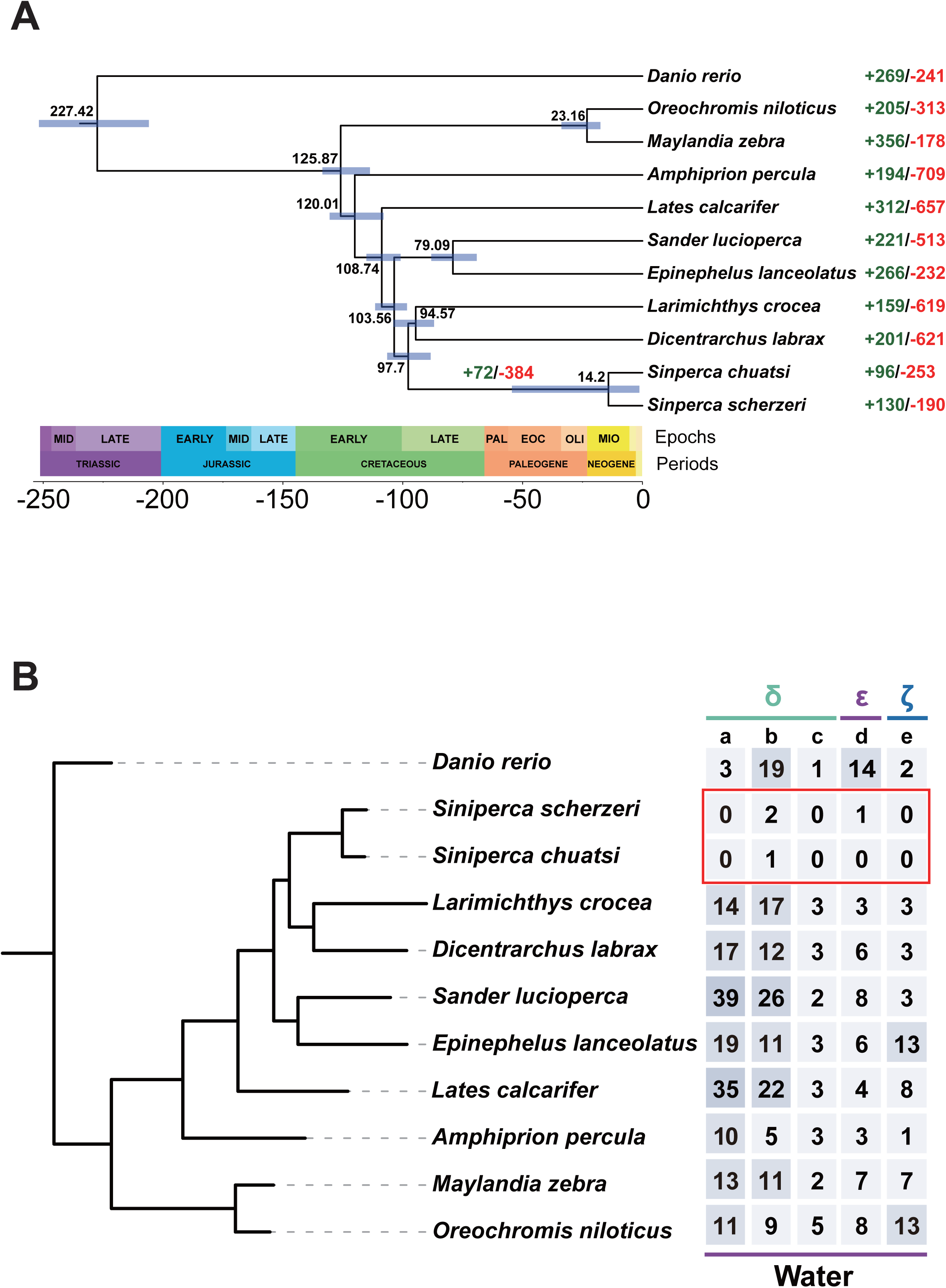
Genetic basis of adaptation in *S. chuatsi* and *S. scherzeri*. **A** A species tree of 10 perciform species with *Danio rerio* as outgroup. The divergence time between species pairs was listed above each node, and 95% confidence interval of the estimated divergence time was denoted as blue bar. The numbers of protein families that were significantly expanded (green) and contracted (red) (P < 0.05) in each species are denoted beside the species names. The numbers of expanded and contracted protein families in the *Siniperca* genus are denoted above the node. **B** OrthoFinder-identified ortholog-groups of olfactory receptor (OR) genes that are contracted in *S. chuatsi* and *S. scherzeri*. a, OG0000155; b, OG0000197; c, OG0000029; d, OG0000041; e, OG0000713.

The olfactory system plays a critical role in feeding, reproduction, predator avoidance and migration of fish [37]. The number and diversity of olfactory receptor (OR) genes, which are positively correlated with the complexity of the olfactory epithelium, contribute to olfactory specialization and ecological adaptation in fish [38]. Sinipercids are extreme piscivore that only accept live prey since the fry start feeding. A previous study showed that *S*. *chuatsi* used vision and mechanoreception but not olfaction for predation [39]. Thus, investigating OR genes can provide insights into the adaptable evolution of olfaction in *S. chuatsi* and *S. scherzeri*. Nine subfamilies of OR genes were classified into two types in vertebrates (Type 1: *α*, *β*, *γ*, *δ*, *ε*, and *ζ*; Type 2: *η*, *θ*, and *κ*), in which *α* and *γ* detect airborne molecules, *δ*, *ε*, *ζ*, and *η* detect water-soluble molecules, and *β* sense both airborne and soluble odorants [40–42]. Interestingly, three OrthoFinder-identified ortholog-groups of OR genes in subfamily δ (OG0000029, OG0000041, OG0000713), one ortholog-group of OR genes in subfamily *ε* (OG0000155), and one ortholog-group of OR genes in subfamily *ζ* (OG0000197) were significantly contracted in the clade of *Siniperca* (**Fig. 3B**). Additionally, a comprehensive genomic screen of functional OR genes found that *S. chuatsi* and *S. scherzeri* had fewer OR genes than seven other perciform fishes (**Supplementary Fig. 6**). These results indicate that the loss of OR genes, especially those in the four ortholog-groups, might be attributed to the special feeding habits of *S. chuatsi* and *S. scherzeri*.

Compared with other perciform fishes, 96 and 130 gene-families were expanded in *S. chuatsi* and *S. scherzeri*, respectively (*P* < 0.05) (**Figure 3A, Supplementary Data 2**). The CREB-binding protein and p300 (CBP/p300) family is a group of transcriptional coactivators that acetylate several histones and non-histone targets [43]. CBP/p300 act as non-DNA-binding co-factors for proteins involved in multiple biological processes, including melanoblast specification, DNA damage response, and circadian rhythm [44–46]. Interestingly, the CBP/p300 family was significantly expanded in the genome of *S. scherzeri* (5 copies) compared with all other perciform fishes (2 copies) (*P* < 0.05) (**Supplementary Fig. 7**). The expansion of CBP/p300 in the genome of *S. scherzeri* may facilitate the quick responses of internal and external stimulates in this species.

Identifying and analyzing PSGs provides insight into how natural selection shapes individual traits during evolution [47]. Therefore, we identified PSGs in the genomes of *S. chuatsi* and *S. scherzeri*. In total, 15 PSGs were identified in the genome of *S. chuatsi* compared with *S. scherzeri* and 8 teleost fishes (*D. rerio*, *Oreochromis niloticus*, *Maylandia zebra*, *Lates calcarifer*, *Larimichthys crocea*, *Dicentrarchus labrax*, *Sander lucioperca*, and *Epinephelus lanceolatus*) (**Supplementary Table 17**). Interestingly, four genes related to growth and development (*smpd3*, *mbtps1*, *fn1a*, *uchl3*) were positively selected in *S. chuatsi*, which may result in the higher growth rate of *S. chuatsi* than that of *S. scherzeri* [48–51]. Additionally, 25 PSGs were found in the *S. scherzeri* genome, compared with *S. chuatsi* and 8 teleost fishes (**Supplementary Table 18**). The selection of three genes that are involved in skin pigmentation (*cdc42*, *rbpjb*, *atrn*) in *S. scherzeri* may contribute to the darker skin color of *S. scherzeri* compared with *S. chuatsi* [52–54].

### The role of SVs in *cis*-regulatory divergence between *S. chuatsi* and *S. scherzeri*

We investigated the role of SVs in *cis*-regulatory divergence between *S. chuatsi* and *S. scherzeri*. The genomic locations of H3K27ac and H3K4me3 were identified in the livers of *S. chuatsi* and *S. scherzeri* using CUT&Tag, respectively (**Supplementary Table 19**). Our analysis identified 17,015 and 15,150 H3K27ac regions (hereafter referred to as peaks), as well as 16,955 and 16,239 H3K4me3 peaks in the genomes of *S. chuatsi* and *S. scherzeri*, respectively (**Supplementary Fig. 8**). The fraction of reads in peaks (FRiP) and transcription start site (TSS) enrichment analyses showed that our CUT&Tag data were of high quality, and sufficient for further analyses (**Supplementary Fig. 9 and Supplementary Table 19**). In addition, we identified open chromatin regions in the genomes of two *Siniperca* fish species using ATAC-seq (**Supplementary Figs. 10 and 11; Supplementary Table 20**). We identified 22,588 nonredundant CREs using CUT&Tag data in the genome of *S. chuatsi*, including 17,623 putative promoters and 4,965 potential distal enhancers, as well as 74,170 open chromatin regions (OCRs) using ATAC-seq data. In addition, 19,858 nonredundant CREs (16,726 putative promoters and 3,132 potential distal enhancers) and 75,972 OCRs were identified in the genome of *S. scherzeri* (**Supplementary Table 21**).

A total of 11,050 differential H3K27ac peaks (|log2FC| >= 1, *P* <= 0.05) between *S. chuatsi* and *S. scherzeri* were identified, of which 6,632 were upregulated and 4,418 were downregulated in *S. chuatsi*. The distribution of differential H3K27ac peaks showed an association with introns and intergenic regions (**Fig. 4A**). We identified 5,437 differential H3K4me3 peaks (|log2FC| >= 1, *P* <= 0.05) between these two species, which were mostly located in introns and promoters (**Fig. 4B**). Additionally, 3,870 differentially expressed genes (|log2FC|LJ>=LJ1, FDRLJ<LJ= 0.05) were identified between the livers of *S. chuatsi* and *S. scherzeri*, of which 2,099 were upregulated and 1,771 were downregulated in *S. chuatsi* compared with *S. scherzeri* (**Supplementary Fig. 12**).

**Figure 4.**
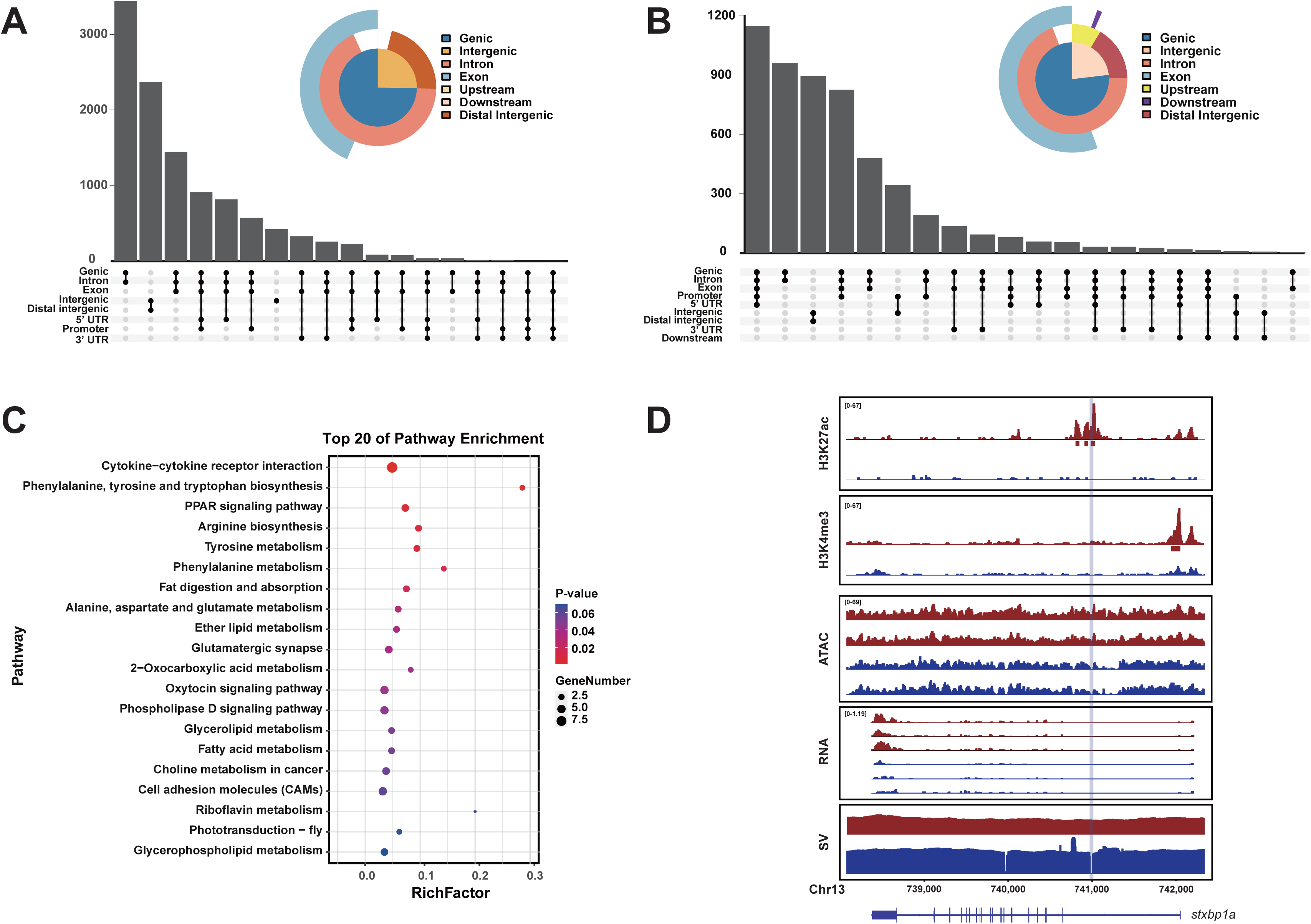
The role of SVs in *cis*-regulatory divergence between *S. chuatsi* and *S. scherzeri*. **A** Genomic distribution of differential H3K27ac peaks. **B** Genomic distribution of differential H3K4me3 peaks. **C** KEGG enrichment of genes that are associated with the differential H3K27ac peaks intersecting the high-confidence SVs. **D** *Cis*-regulatory divergence between *S. chuatsi* (Red) and *S. scherzeri* (Blue) at *stxbp1a* gene. The intensity of H3K27ac, H3K4me3, ATAC-seq, as well as gene expression level are shown. In addition, mapping coverages of Nanopore reads are plotted to indicate genomic structural variants. A deletion in *S. scherzeri* is denoted with a light blue shade. MACS2 identified H3K27ac and H3K4me3 peaks are denoted with rectangles below tracks. Transcripts (with exons as boxes) are depicted.

Based on the high-quality genome assemblies, SVs were identified between the genomes of *S. chuatsi* and *S. scherzeri* using three different tools (**Supplementary Fig. 13**). SyRI and Smartie-SV identify SVs by aligning two genome assemblies, while NGMLR-Sniffles calls SVs by aligning Nanopore reads to reference genome sequences [55–57]. By merging the results of these three tools, 75,988 deletions, 84,189 insertions, 1,714 duplications, and 310 inversions were detected between the genomes of *S. chuatsi* and *S. scherzeri*.

A total of 1,358 SVs overlapped with differential H3K27ac peaks between the genomes of *S. chuatsi* and *S. scherzeri*. We validated these SVs using Samplot and obtained 905 high-confidence SVs [58]. In total, 1,205 genes were associated with the differential H3K27ac peaks intersecting high-confidence SVs. In addition, 301 out of the 1,205 genes showed significantly differential expression (|log2FC| >= 1, FDR < 0.05) between *S. chuatsi* and *S. scherzeri*. Kyoto Encyclopedia of Genes and Genomes (KEGG) enrichment analysis showed that the 301 DEGs were mainly enriched in pathways related to lipid and amino acid metabolism, including arginine biosynthesis (ko00220), fat digestion and absorption (ko04975), and fatty acid metabolism (ko01212) (**Fig. 4C**). Interestingly, three genes involved in fatty acid catabolism (*acsl1a*, *acadl*, *got2a*) showed a higher signal intensity of H3K27ac peaks and gene expression levels in *S. chuatsi* than in *S. scherzeri* (**Supplementary Figs. 14-16**). The expression of *acsl1a*, one of the key members of the long-chain acyl-CoA synthetase family responsible for fatty acid degradation and lipid synthesis, is positively correlated with lipid uptake [59, 60]. The *acadl* gene plays a critical role in lipid catabolism by catalyzing the initial step for β-oxidation of long-chain fatty acyl-CoAs [61]. *got2a* promotes fatty acid metabolism by transporting long-chain fatty acids into the cell [62]. The elevated expression of these three genes suggests that *S. chuatsi* can utilize fatty acids more efficiently than *S. scherzeri*, which may contribute to the higher growth rate in *S. chuatsi* compared with *S. scherzeri*.

Three genes (*stxbp1a*, *mkln1*, *myca*) involved in skin pigmentation showed differential gene expression levels and H3K27ac peak intensities between *S. chuatsi* and *S. scherzeri*. A deletion in the first intron of *stxbp1a* completely removes a putative enhancer in *S. scherzeri*, and the expression of this gene is lower in *S. scherzeri* than in *S. chuatsi* (**Fig. 4D**). *stxbp1a* mutant zebrafish have darker pigmentation on their heads and backs [63]. Therefore, reduced expression of this gene in *S. scherzeri* may lead to darker pigmentation in this species. A deletion in the first intron of *mkln1* reduces the intensity of H3K27ac peaks in *S. chuatsi*, resulting in lower expression level of this genes in *S. chuatsi* than in *S. scherzeri* (**Supplementary Fig. 17**). *mkln1* mutant mice develop brighter fur over time [64]. Thus, darker pigmentation in *S. scherzeri* may be attributed to elevated expression of *mkln1*. These results indicate that *cis*-regulatory variation caused by SVs results in expression divergence of these two genes, which in turn leads to pigmentation differences between *S. chuatsi* and *S. scherzeri*.

In addition to genes related to metabolism and skin pigmentation, we found the intensities of H3K27ac peaks associated with four immune genes (*prss16*, *lama4*, *cd22*, *tecpr1b*) were significantly decreased (|log2FC| >= 1, *P* <= 0.05) in *S. chuatsi* compared with *S. scherzeri* (**Supplementary Table 22**). Moreover, the expression levels of these genes in spleen, one of the major immune organs in fish, are substantially higher in *S. scherzeri* than in *S. chuatsi*. The thymus-specific serine protease (TSSP), which is encoded by *prss16*, plays an critical role in T cell maturation [65]. The ability of immune cells to penetrate the vessel wall is reduced in *lama4* mutant mice [66]. The *cd22* gene is a regulator of innate and adaptive B cell responses [67]. And a *tecpr1*-dependent pathway is important in targeting bacterial pathogens for selective autophagy [68]. Previous studies showed that *S. scherzeri* was more resistant to disease than *S. chuatsi* [25]. Therefore, the *cis*-regulatory variation caused by SVs in these four gene may lead to divergence in disease resistance between *S. scherzeri* and *S. chuatsi*.

### Divergence of Broad H3K4me3 domains between *S. chuatsi* and *S. scherzeri*

To determine whether broad H3K4me3 domains are involved in phenotypic divergence between *S. chuatsi* and *S. scherzeri*, we identified and compared the broad H3K4me3 peaks in the genomes of these two species. The results from our analysis revealed 491 and 481 broad H3K4me3 peaks (top 3% broadest H3K4me3 domains) in the genomes of *S. chuatsi* and *S. scherzeri*, respectively. The broad H3K4me3 peaks were mostly found close to genes, extending both the 5’ and 3’ of TSSs (**Figure 5A and B**). Consistent with the previous results, the signal intensity of broad H3K4me3 peaks is lower than that of narrow H3K4me3 peaks [20]. In addition, 93.5% and 92.3% of the broad H3K4me3 peaks in *S. chuatsi* and *S. scherzeri* overlapped with H3K27ac peaks, respectively, corroborating the view that broad H3K4me3 domains tend to overlap with H3K27ac domains (**Supplementary Table 23**) [69]. A total of 194 differential broad H3K4me3 peaks (|log2FC| >= 1, *P* <= 0.05) were identified between the genomes of *S. chuatsi* and *S. scherzeri*, of which 119 were upregulated in *S. chuatsi*, and 75 were downregulated in *S. chuatsi*. We used KEGG pathway analysis to characterize enriched functions for the genes associated with differential broad H3K4me3 peaks (**Fig. 5C; Supplementary Table 24**). It is noteworthy that genes associated with differential broad H3K4me3 peaks are enriched in several pathways related to cancer, including microRNA in cancer (ko05206), pathways in cancer (ko05200), and choline metabolism in cancer (ko05231). Among the genes enriched in pathways in cancer, the gene expression level (log2FC >= 1, FDR <= 0.05) as well as the intensity of broad H3K4me3 peaks (log2FC >= 1, *P* <= 0.05) of 4 genes (*ccnd2a*, *egln2*, *kita*, *f2r*) were significantly upregulated in *S. chuatsi* compared with *S. scherzeri* (**Fig. 5D; Supplementary Figs. 18-20; Supplementary Table 25**). Most of these genes are involved in various developmental processes, including cardiovascular development and melanocyte development [70–72]. The results indicate that the divergence of broad H3K4me3 domains contribute to the phenotypic divergence between *S. chuatsi* and *S. scherzeri*.

**Figure 5.**
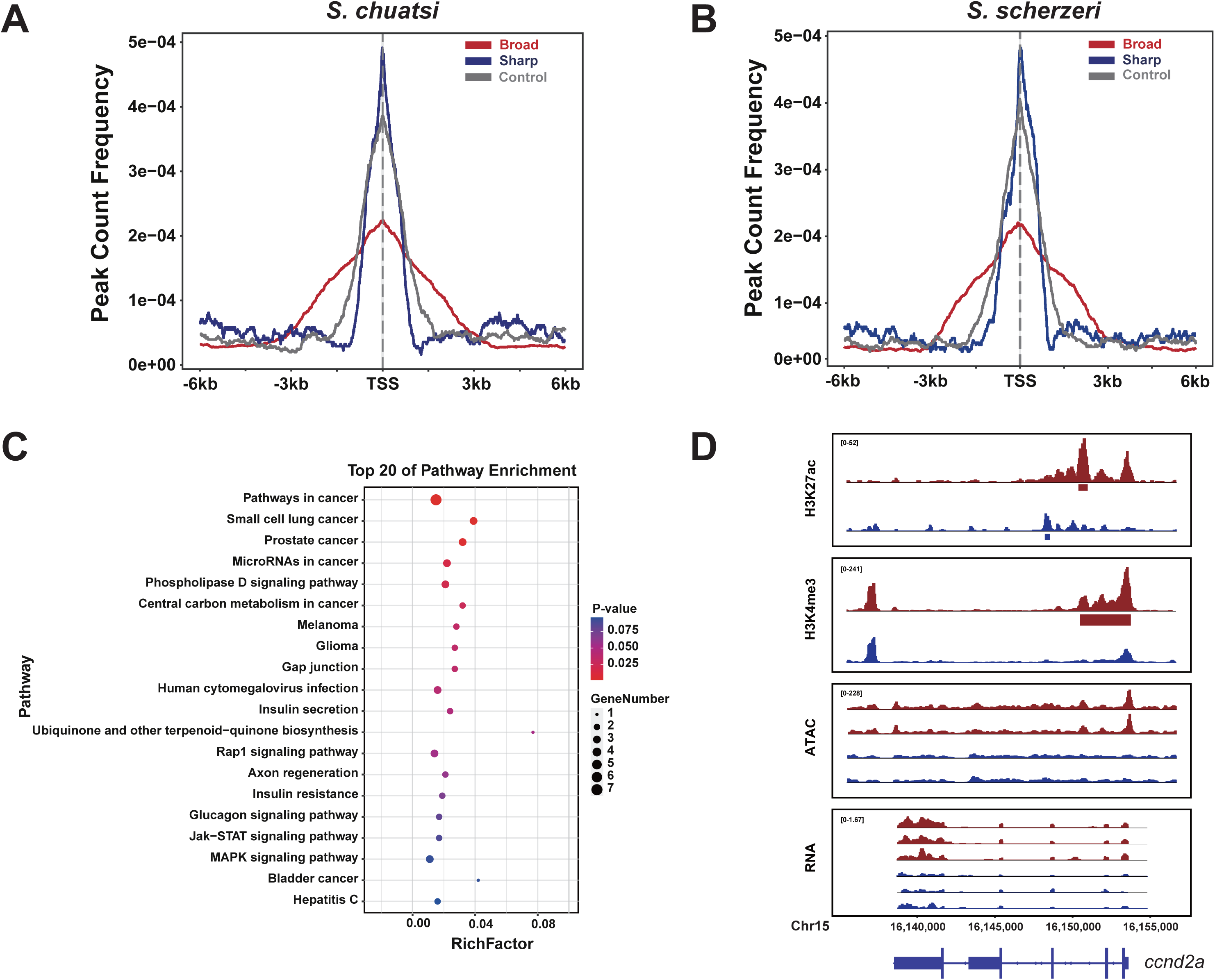
Divergence of Broad H3K4me3 domains between *S. chuatsi* and *S. scherzeri*. Signal intensities of 400 broad H3K4me3 peaks (red), 400 sharp H3K4me3 peaks (blue), as well as 400 randomly selected H3K4me3 peaks (control, grey) of *S. chuatsi* (**A**) and *S. scherzeri* (**B**). **C** KEGG enrichment of genes that are associated with divergent broad H3K4me3 peaks. **D** *Cis*-regulatory divergence between *S. chuatsi* (Red) and *S. scherzeri* (Blue) at *ccnd2* gene. The intensity of H3K27ac, H3K4me3, ATAC-seq, as well as gene expression level are shown. MACS2 identified H3K27ac and broad H3K4me3 peaks are denoted with rectangles below tracks. Transcripts (with exons as boxes) are depicted.

## Discussion

By integrating Nanopore and Hi-C sequencing strategies, we generated two high-quality, chromosome-level assemblies of two closely related *Siniperca* fish species, *S. chuatsi* and *S. scherzeri*. Based on these two nearly complete genome assemblies, we first studied the evolutionary history of these two species. Second, we dissected the role of SVs intersecting *cis*-regulatory elements in phenotypic divergence between *S. chuatsi* and *S. scherzeri*. Finally, the role of broad H3K4me3 domains in phenotypic divergence between these two closely related species was investigated.

### Demographic history analysis of *S. chuatsi* and *S. scherzeri* provides insights into the evolution of sinipercid fish

*Siniperca chuatsi* and *S. scherzeri* are the two most widely distributed sinipercids, indicating that they are well-adapted to diverse environments in East Asia, especially in China [22]. In addition, *S. chuatsi* is one of the major aquaculture fish species in China due to its excellent flesh quality [73]. Thus, investigating the evolutionary history of these two species provides insights into the adaptation in this subfamily and facilitates breeding of *S. chuatsi*. The ancestors of sinipercid fish repeatedly invaded the freshwater system from the ocean in the early Cenozoic Era [74]. The common ancestors of subfamily Sinipercidae and genus *Siniperca* are estimated to occur 43.4 Ma and 12.2 Ma, respectively [22]. However, the demographic history of sinipercid fish is largely unknown. We estimated the histories of ancestral population sizes of *S*. *chuatsi* and *S. scherzeri* using two approaches. PSMC inferred that the ancestral population size of *S*. *chuatsi* declined during the early phase of the Mid-Pleistocene Transition (~0.9 Ma), where the duration of the Pleistocene glacial cycles increased from 41 to 100 ka [31]. SMC++ analysis revealed a population bottleneck at the beginning of the last glacial period (~90 ka), after the Eemian interglacial period (129~116 ka). These results suggest that the colonization of new habitats and diversification in *S*. *chuatsi* are largely affected by the changes in temperature.

Sinipercids disperse over two major zoogeographic regions (Palearctic and Oriental) in China. Moreover, two populations of *S. scherzeri* were identified in northern and southern China [75]. Therefore, we inferred the dynamic changes in the ancestral population sizes of two *S. scherzeri* populations from northern (Jinlin Province) and southern (Guangdong Province) China. SMC++ inferred that *S. scherzeri* from South China have experienced a decline in effective population size at ~400 ka, after the Mid-Brunhes Event (~430 ka), while the ancestral population size of *S. scherzeri* from North China declined at ~300 ka. These results imply that different populations of this widely distributed species have distinct histories of local adaptation.

### *Cis*-regulatory divergence caused by SVs plays an important role in phenotypic divergence between *S. chuatsi* and *S. scherzeri*

Gene expression plays a major role in the form, fitness, and function of organisms [76]. It has been shown that differences in *cis*-regulatory activity contribute to gene expression divergence over a wide range of evolutionary divergence scales and play a major role in phenotypic divergence [77–79]. SVs are considered to be essential in the phenotypic evolution of both animals and plants [80]. Additionally, SVs have been found to alter the *cis*-regulatory activity of several developmental regulatory genes, resulting in morphological divergence between species [3]. However, most of the SVs identified using short-read sequencing are unreliable, leaving a comprehensive investigation of the impacts of SVs on *cis*-regulatory divergence largely impossible. With the advent of third-generation sequencing technologies, researchers can reveal the role of SVs in phenotypic variation among diverse species. We identified genome-wide high-quality SVs between *S*. *chuatsi* and *S. scherzeri* and investigated their role in the divergence of CREs marked with H3K27ac modification. Differentially expressed genes associated with these SV-intersected CREs were mainly enriched in pathways related to lipid and amino acid metabolism. We found an elevated signal intensity of H3K27ac peaks and expression levels of several genes involved in amino acid metabolism and fatty acid catabolism in *S*. *chuatsi*, indicating that the efficiency of lipid and amino acid metabolism is higher in *S*. *chuatsi* than in *S. scherzeri*. The growth rate of *S*. *chuatsi* is substantially higher than that of *S. scherzeri* [25]. Organisms with high growth rates tend to have higher rates of lipid and amino acid metabolism [81]. These results suggest that *cis*-regulatory divergence induced by SVs causes expression divergence of genes involved in lipid and amino acid metabolism, which in turn leads to differences of growth rates between *S*. *chuatsi* and *S. scherzeri*. Additionally, we found that SVs caused *cis*-regulatory divergence in genes involved in skin pigmentation between *S*. *chuatsi* and *S. scherzeri*. The resulting gene expression divergence leads to darker pigmentation in *S. scherzeri*. Lastly, the expression levels of four immune genes associated with SV-intersected divergent H3K27ac peaks are substantially higher in *S. scherzeri* than in *S*. *chuatsi*. These four genes play critical role in organism’s immunity. This suggests that SVs caused *cis*-regulatory variation contributes to divergence in disease resistance between *S*. *chuatsi* and *S. scherzeri*. Taken together, our results suggest that *cis*-regulatory divergence caused by SVs play an important role in phenotypic divergence between two closely related *Siniperca* species.

The relative contribution of coding and regulatory changes in speciation has been under debate. Studies in diverse organisms showed coding sequence variations of adaptive loci contributing to their speciation and adaptation [82]. The contrasting hypothesis suggests that modifications of gene expression by changes in regulatory regions play a prominent role in evolution and adaptation [83]. Our analysis of PSGs using protein coding sequences found that three genes related to pigmentation were positively selected in *S. scherzeri* compared with *S. chuatsi* and 8 teleost fishes. Additionally, *cis*-regulatory divergence was found in two pigmentation genes between *S. chuatsi* and *S. scherzeri*. These results indicate that both coding and regulatory changes contribute to speciation and adaptation in *S. chuatsi* and *S. scherzeri*.

### Broad H3K4me3 domains associated with cancer-related genes in *S. chuatsi* and *S. scherzeri* and contribute to their phenotypic divergence

There are two classes of H3K4me3 domains, including the intensively studied narrow domain and the currently identified broad domain. Most of H3K4me3-enriched nucleosomes are detected as narrow peaks (1~2 kb), which are associated with the promoters of actively transcribed genes and serve as a switch of gene transcription [17]. A small number of broad H3K4me3 domains, which can span up to 60 kb, were recently identified in several species including mammals, flies, worms, and plants [69]. Broad H3K4me3 domains are correlated with increased transcription elongation and enhancer activity, resulting in high expression levels of associated genes [20]. This special histone modification preferentially marks genes associated with cell identity and cell-specific function [21]. A study revealed that broad H3K4me3 domains modulated maternal-to-zygotic transition in mouse oocytes [19]. Furthermore, broad H3K4me3 domains were found to explicitly mark cancer-suppressor genes in normal human cells [20]. This suggests that broad H3K4me3 domains may play a role in phenotypic divergence in animals by regulating developmental processes. However, comparative analysis of broad H3K4me3 domains between closely related species is still lacking. Studies of this domain are largely restricted to a single species or distantly related species. In this study, we compared broad H3K4me3 domains in the genomes of two closely related fish species. Genes that are associated with the divergent domains are mostly enriched in cancer-related pathways. Moreover, most of these genes are involved in various developmental processes. These results suggest that the association of broad H3K4me3 domains and cancer-related genes has an ancient origin. The divergence of broad H3K4me3 domains contributes to phenotypic divergence between two closely related species.

## Conclusions

In conclusion, we generated high-quality chromosome-level genome assemblies of *S. chuatsi* and *S. scherzeri*, facilitating comparative genomic analysis of these two closely related fish species. Firstly, demographic analysis revealed that *S. chuatsi* had experienced two population bottlenecks in the early phase of the Mid-Pleistocene Transition (~0.9 Ma) and at the beginning of the last glacial period (~90 ka), respectively. This suggests the colonization of new habitats and diversification in *S*. *chuatsi* are largely affected by the changes in temperature. *S. scherzeri* from northern China have experienced a decline in effective population size ~100 ka later than *S. scherzeri* from southern China, indicating that different populations of this widely distributed species have distinct histories of local adaptation. Secondly, based on the chromosome-level assemblies, we identified high-quality SVs between the genomes of *S. chuatsi* and *S. scherzeri*. Our integrative analysis of SVs and H3K27ac domains found *cis*-regulatory divergence caused by SVs play an essential role in the divergence of lipid and amino acid metabolism, skin pigmentation, and immunity between *S. chuatsi* and *S. scherzeri*. This suggests *cis*-regulatory divergence caused by SVs play an important role in phenotypic divergence between these two closely related species. Finally, we found that the divergence of broad H3K4me3 domains contributed to phenotypic divergence between *S. chuatsi* and *S. scherzeri*. And the association of broad H3K4me3 domains and cancer-related genes has an ancient origin.

## Materials and Methods

### Genome sequencing

One male *S. chuatsi* individual and one *S. scherzeri* male individual, which were collected from Fujian province, China, were used for genome sequencing. All operations of fishes were approved by the Institutional Animal Care and Use Committee of Sun Yatsen University (Protocol number SYSU-IACUC-2020-B0975). High-quality DNA was extracted from muscle cells of *S. chuatsi* and *S. scherzeri* using DNeasy Blood & Tissue Kits (Qiagen) in accordance with the manufacturer’s protocol. Quality and quantity of the DNA were measured via standard agarose-gel electrophoresis and with a Qubit 3.0 Fluorometer (Invitrogen). Nanopore sequencing libraries of *S. chuatsi* and *S. scherzeri* were constructed and sequenced by Nanopore PromethION platform (Oxford Nanopore Technologies), respectively (140X raw-read coverage for *S. chuatsi*; 127X raw-read coverage for *S. scherzeri*). For Illumina sequencing, short-insert paired-end (PE) (150 bp) DNA libraries of *S. chuatsi* and *S. scherzeri* were constructed in accordance with the manufacturer’s instruction, respectively. Sequencing for the PE libraries were performed with 2X150 bp chemistry on the Illumina NovaSeq 6000 platform (Illumina).

### Transcriptome sequencing for genome annotation

Samples from eye, gill, heart, intestine, kidney, stomach, testis, liver, and spleen were collected for *S. chuatsi* and *S. scherzeri* to construct sequencing libraries of strand-specific RNA-sequencing (RNA-seq). Total RNA was extracted with TRIzol reagent (Invitrogen). The purity and integrity were determined with NanoDrop 2000 spectrophotometer (Thermo Fisher Scientific) and Bioanalyzer 2100 system (Agilent). The mRNA was enriched from total RNA using poly-T oligo-attached magnetic beads. And rRNA was removed using TruSeq Stranded Total RNA Library Prep kit (Illumina). Paired-end library was constructed using the VAHTSTM mRNA-seq V2 Library Prep Kit for Illumina (Vazyme) and sequenced with 2×150 bp chemistry on the Illumina HiSeq NovaSeq 6000 platform (Illumina).

To construct full-length RNA-seq libraries, total RNA was extracted from muscle cells of *S. chuatsi* and *S. scherzeri* with TRNzol Universal reagent (TIANGEN), respectively. The integrity of the RNA was determined with NanoDrop One Microvolume UV-Vis spectrophotometer (Thermo Fisher Scientific) and Bioanalyzer 2100 system (Agilent Technologies). The concentration of the RNA was determined with Qubit 3.0 Fluorometer (Invitrogen). The SMARTer PCR cDNA Synthesis Kit (Takara) was used for first-strand cDNA synthesis. PCR was performed to generate double-stranded cDNA using PrimeSTAR GXL DNA polymerase (Takara). Sequencing libraries of both species were then constructed using SMRTbell Express Template Prep Kit 2.0 and Sequel Binding Kit 3.0 (Pacific Biosciences) in accordance with the manufacturer’s protocol. Sequencing was performed on a PacBio Sequel II platform (Pacific Bioscience).

### Genome size estimation and genome assembly

Low-quality (reads with ≥10% unidentified nucleotide and/or ≥ 50% nucleotides having phred score < 5) and sequencing-adapter-contaminated Illumina reads were filtered and trimmed with Fastp (v0.21.0) [84]. The resulted high-quality Illumina reads were used in the following analyses. The sizes and heterozygosity of *S. chuatsi* and *S. scherzeri* genomes were estimated using high-quality Illumina reads by *k*-mer frequency-distribution method, respectively. The number of *k*-mers and the peak depth of *k*-mer sizes at 21 was obtained using Jellyfish (v2.3.0) [85] with the -*C* setting. The results of Jellyfish were then inputted into GenomeScope2 (v1.0.0) [86] to estimate the genome size and heterozygosity rate.

Low-quality Nanopore reads were filtered using previous published Python script [87]. The filtered reads were then corrected using NextDenovo (v1.0) (https://github.com/Nextomics/NextDenovo). Draft-genome assemblies for *S. chuatsi* and *S. scherzeri* were generated using filtered and corrected reads with WTDBG (v1.2.8) [27], respectively. The contigs of the two draft assemblies were subject to error correction using high-quality Illumina reads with Pilon (v1.23) [88] three times, respectively.

We used Hi-C to correct misjoins, to order and orient contigs, and to merge overlaps. Liver samples of *S. chuatsi* and *S. scherzeri* were collected to construct Hi-C libraries. Hi-C libraries were constructed using the previously published approach [89]. Hi-C libraries of *S. chuatsi* and *S. scherzeri* were sequenced with 2×150 bp chemistry on the Illumina MiSeq platform (Illumina), respectively. Low-quality sequencing reads were filtered using fastp (v0.21.0) [84] with default parameters. Filtered Illumina reads were aligned to the respective assembled contigs of each species using Juicer (v1.5.7) [90]. Scaffolding was accomplished with a 3D-DNA pipeline (v180419) [91]. Juicebox (v1.9.9) was used to modify the order and directions of some scaffolds in a Hi-C contact map and to help in the determination of chromosome boundaries [92].

The completeness and quality of the final assemblies of *S. chuatsi* and *S. scherzeri* were first evaluated using Benchmarking Universal Single-Copy Orthologs (BUSCO) (v4.0.5) [93] against the conserved Eukaryota dataset (odb10). Second, RNA-seq reads of brain, intestine, liver, and muscle of *S. chuatsi* were downloaded from NCBI [28]. Previously published and our RNA-seq reads of *S. chuatsi* were both aligned to previously published and our assemblies of *S. chuatsi* genome using HISAT2 (v2-2.1) [94], respectively. RNA-seq reads of *S. scherzeri* generated in this study were aligned to previously published and our assemblies of *S. scherzeri* genome using HISAT2 (v2-2.1), respectively. Third, Merqury (v1.3) [29] was used to assess the completeness and quality of four assemblies with *k*-mer set to 20. Fourth, Nanopore reads of *S. chuatsi* were aligned to previously published and our assemblies of *S. chuatsi* genome using Minimap2 (v2.16-r922) [95], respectively. And Nanopore reads of *S. scherzeri* were aligned to previously published and our assemblies of *S. scherzeri* genome using Minimap2 (v2.16-r922), respectively. The alignment results of some genomic regions were visualized using Samplot (v1.3.0) [58]. Lastly, our assemblies of *S. chuatsi* and *S. scherzeri* genomes were aligned to the respective published assemblies using Minimap2 (v2.16-r922). The results were visualized using the script pafCoordsDotPlotly.R (https://github.com/tpoorten/dotPlotly).

### Genome annotation

Repetitive elements in the assembly were identified by *de novo* predictions using RepeatMasker (v4.1.0) (https://www.repeatmasker.org/). RepeatModeler (v2.0.1) [96] was used to build the *de novo* repeat libraries of *S. chuatsi* and *S. scherzeri*. To identify repetitive elements, sequences from the *S. chuatsi* and *S. scherzeri* assemblies were aligned to the *de novo* repeat library using RepeatMasker (v4.1.0), respectively. Additionally, repetitive elements in *S. chuatsi* and *S. scherzeri* genome assemblies were identified by homology searches against known repeat databases using RepeatMasker (v4.1.0).

Protein-coding genes in *S. chuatsi* and *S. scherzeri* genomes were predicted with three approaches: homology-based prediction, *ab initio* prediction and RNA-seq-based prediction. For homology-based prediction, protein-coding sequences of *Danio rerio*, *Gasterosteus aculeatus*, *Takifugu rubripes*, *Oryzias latipes* and *Tetraodon nigroviridi* were downloaded from Ensembl (v96) [97], and aligned to *S. chuatsi* and *S. scherzeri* genomes using tblastn, respectively. GenomeThreader (v1.7.0) [98] was employed to predict gene models based on the alignment results with an E-value cut-off of 10^−5^. For *ab initio* gene prediction, the short-read RNA-seq of the 9 samples (eye, gill, heart, intestine, kidney, stomach, testis, liver, spleen) of *S. chuatsi* and *S. scherzeri* were aligned to the respective assembled genome sequences using STAR (v2.7.0) [99]. Additionally, full-length RNA-seq reads of *S. chuatsi* and *S. scherzeri* was aligned to the respective assembled genome sequences using GMAP (2018-07-04) [100]. Gene models were predicted based on the alignment results of short-read and full-length RNA-seq reads using BRAKER2 (v2.1.5) [101].

For RNA-seq-based prediction, the short-read RNA-seq reads of *S. chuatsi* and *S. scherzeri* were first aligned to respective reference sequences using HISAT2 (v2-2.1). Gene models were predicted based on the alignment results of HISAT2 using StringTie (v2.1.4) [102], and coding regions were identified using TransDecoder (v5.5.0). Second, short-read RNA-seq reads of *S. chuatsi* and *S. scherzeri* were assembled using Trinity (v2.8.5) [103], respectively. Third, full-length RNA-seq reads of *S. chuatsi* and *S. scherzeri*were assembled using Iso-Seq3 (v3.1) (https://github.com/PacificBiosciences/IsoSeq), respectively. Iso-Seq3 assembled full-length RNA-seq reads were subject to error correction using high-quality Illumina reads with LoRDEC (v0.5.3) [104]. Finally, Program to Assemble Spliced Alignments (PASA) (v2.5.0) were used to predict gene models in genomes of both species based on the assembly results of Trinity and Iso-Seq3 with StringTie predicted gene models as a reference. Coding regions of PASA predicted gene models were then identified using TransDecoder (v5.5.0) [103].

Gene models of *S. chuatsi* and *S. scherzeri* predicted by BRAKER2, GenomeThreader, and PASA were integrated into a nonredundant consensus-gene set using EVidenceModeler (v1.1.1) [105], respectively. EVidenceModeler integrated gene models were updated with short-read and full-length RNA-seq reads using PASA (v2.5.0) three times. The completeness of predicted gene models of the two species were evaluated using Benchmarking Universal Single-Copy Orthologs (BUSCO) (v4.0.5) [93] against the conserved Vertebrata dataset (odb10).

To assign functions to the predicted proteins, we aligned the *S. chuatsi* and *S. scherzeri* protein models against NCBI nonredundant (NR) amino acid sequences, UniProt, Translated EMBL-Bank (trEMBL), Cluster of Orthologous Groups for eukaryotic complete genomes (KOG), SwissProt database using blastp with an E-value cutoff of 10^−5^. Protein models were also aligned against eggNOG database [106] using eggNOG-Mapper [107]. Finally, KEGG annotation of the protein models was performed using BlastKOALA [108].

### Genome resequencing and SNP calling

To investigate the evolutionary history and adaptation of *S. chuatsi* and *S. scherzeri*, we collected 6 wild *S. chuatsi* individuals from Hunan province, China, 6 wild *S. scherzeri* individuals from Guangdong province, China, and 6 wild *S. scherzeri* individuals from Jilin province, China. Sequencing libraries were constructed following Illumina TruSeq Nano DNA HT sample preparation kit in accordance with the manufacturer’s protocol. All individuals were whole-genome re-sequenced to an average coverage of 12.5× with 2×150 bp chemistry on the Illumina NovaSeq 6000 platform (**Supplementary Table 16**).

Low-quality and sequencing-adapter-contaminated Illumina reads were filtered and trimmed with Trimommatic (v0.36) [109]. High-quality pair-end reads of *S. chuatsi* and *S. scherzeri* were aligned to the respective reference sequences using BWA (v0.7.17) with “mem” function [110]. PCR duplicates were removed using MarkDuplicate program of Picard (v2.18.27) (https://broadinstitute.github.io/picard/). SNP variants were identified using HaplotypeCaller program of the Genome Analysis Toolkit (GATK) (v4.1.0.0). Raw SNP calling dataset were filtered using VariantFiltration program of GATK (v4.1.0.0) with parameters “QDLJ<LJ2.0 || QUAL < 30.0 || FSLJ>LJ60.0 || MQLJ<LJ40.0 || MQRankSumLJ<LJ−LJ12.5 || ReadPosRankSumLJ<LJ−LJ8.0 || SORLJ>LJ3.0”.

### Demographic inference of *S. chuatsi* and *S. scherzeri*

Historical effective population sizes of *S. chuatsi* and *S. scherzeri* were inferred using both SMC++ and PSMC. Historical effective population sizes of *S. chuatsi* and *S. scherzeri* were first estimated using SNP variants of 6 wild-caught *S. chuatsi* individuals and 6 wild-caught *S. scherzeri* individuals using SMC++ (v1.15.3). The substitution mutation rate and generation time of *S. chuatsi* and *S. scherzeri* was set to 2.22 × 10^-9^ and 2 years according to the previous study of *Siniperca knerii* (Big-eye Mandarin Fish) [111]. High-quality Illumina reads of *S. chuatsi* and *S. scherzeri* generated for genome assembly were aligned to the respective reference sequences using BWA (v0.7.17) with “mem” function. Genetic variants were identified using Samtools (v1.9-52) [112]. Whole-genome consensus sequence was generated using the genetic variants using Samtools (v1.9-52) [112]. PSMC (v0.6.5) [30] was used to infer population size history of *S. chuatsi* and *S. scherzeri* using the whole genome consensus sequences. The substitution mutation rate and generation time of *S. chuatsi* and *S. scherzeri* was set to 2.22 × 10^-9^ and 2 years according to the previous study of *S. knerii* [111].

### Phylogenetic reconstruction

Protein sequences of 8 species (*D. rerio*, *O. niloticus, M. zebra*, *Amphiprion ocellaris*, *L. calcarifer*, *L. crocea*, *D. labrax*, *Sander lucioperca*) were downloaded from Ensembl (v96) [97]. Protein sequences of *E. lanceolatus* was downloaded from NCBI [113]. Protein sequences shorter than 50 amino acids were removed. OrthoFinder (v 2.5.4) [114] was applied to determine and cluster gene families among these 9 species as well as *S. chuatsi* and *S. scherzeri*. Gene clusters with >100 gene copies in one or more species were removed. Single-copy orthologs in each gene cluster were aligned using MAFFT (v7.490) [115]. The alignments were then trimmed using Gblocks (v0.91b) [116]. The phylogenetic tree was reconstructed with the trimmed alignments using a maximum-likelihood method implemented in IQ-TREE2 (v2.2.0) with *Danio rerio* as outgroup. The best-fit substitution model was selected by using ModelFinder algorithm [117]. Branch supports were assessed using the ultrafast bootstrap (UFBoot) approach with 1000 replicates [118].

Divergence time was estimated using MCMCtree module of the PAML package (v4.9) [119]. MCMCtree analysis was performed using the maximum-likelihood tree that was reconstructed by IQ-TREE2 as a guide tree and calibrated with the divergent time obtained from TimeTree database (minimum = 206 million years and soft maximum = 252 million years between *D. rerio* and *O. niloticus*; minimum = 18.08 million years and soft maximum = 35.16 million years between *M. zebra* and *O. niloticus*; minimum = 82 million years and soft maximum = 131 million years between *A. percula* and *O. niloticus*; minimum = 104 million years and soft maximum = 145 million years between *A. percula* and *L. calcarifer*; minimum = 94 million years and soft maximum = 115 million years between *S. lucioperca* and *L. calcarifer*; minimum = 69 million years and soft maximum = 88 million years between *S. lucioperca* and *E. lanceolatus*; minimum = 99 million years and soft maximum = 127 million years between *S. lucioperca* and *L. crocea*; minimum = 87 million years and soft maximum = 105 million years between *D. labrax* and *L. crocea*) (retrieved June 2021) [120].

### Gene family expansion and contraction analysis

CAFÉ (v5) [121] was applied to determine the significance of gene-family expansion and contraction among 11 teleost species based on the MCMCtree generated ultrametric tree and OrthoMCL determined gene clusters that were used for reconstructing species tree.

A phylogenetic tree was reconstructed to investigate the evolutionary relationships of EP300 genes from *S. chuatsi*, *S. scherzeri*, and other teleost species. Protein sequences of EP300 from *D. rerio*, *O. niloticus, M. zebra*, *L. calcarifer*, *L. crocea*, *D. labrax*, *S. lucioperca*, *E. lanceolatus*, *G. aculeatus* were downloaded from Ensembl (v96) [97]. The protein sequences were aligned using MAFFT (v7.490) [115]. The phylogenetic tree was reconstructed with the alignments using a maximum-likelihood method implemented in IQ-TREE2 (v2.2.0) with an EP300 protein sequence (XP_040032248.1) from *G. aculeatus* as outgroup. The best-fit substitution model was selected by using ModelFinder algorithm [117]. Branch supports were assessed using the ultrafast bootstrap (UFBoot) approach with 1000 replicates [118].

### Analysis of Olfactory Receptor Genes

Olfactory receptor (OR) genes in the genomes of *D. rerio*, *O. niloticus*, *M. zebra*, *A. ocellaris*, *L. calcarifer*, *L. crocea*, *D. labrax*, *S. lucioperca*, *E. lanceolatus*, *S. chuatsi*, and *S. scherzeri* were identified using a previously described approach [122]. In brief, tblastn search was performed with a E-value cut-off of 10^-10^ using a set of known functional OR genes from *D. rerio*, *L. crocea*, *L. calcarifer*, *O. niloticus*, *O. latipes*, *T. rubripes* as queries [123]. The best-hit regions were extracted and extended 5000 bp in both 3’ and 5’ direction along the genome sequences using Samtools (v1.9-52) [112]. The structures of genes were predicted using the extended best-hit regions with Exonerate (v2.4.0) [124]. The resulted protein-coding sequences were aligned against UniProt database using blastp (v2.8.1) and the ones that have the highest alignment scores to known ORs were retained. We used CD-HIT (v4.6.2) [125] to remove redundant sequences and cluster filtered OR sequences. We classified OR genes with an intact sequence and without loss-of-function mutation as functional genes. The functional OR protein sequences were aligned using MAFFT (v7.490) [115]. The phylogenetic tree was reconstructed with the alignments using a maximum-likelihood method implemented in IQ-TREE2 (v2.2.0). The best-fit substitution model was selected by using ModelFinder algorithm [117]. Branch supports were assessed using the ultrafast bootstrap (UFBoot) approach with 1000 replicates [118].

### Population genetic analyses

To determine genetic differentiation among wild-caught *S. chuatsi* and *S. scherzeri* populations, principal component analysis (PCA) was performed using genome-wide SNPs of 6 *S. chuatsi* individuals from Hunan province, China, 6 wild *S. scherzeri* individuals from Guangdong province, China, and 6 wild *S. scherzeri* individuals from Jilin province, China with plink (v1.90) [126]. Genome-wide nucleotide diversity (π) of each population was calculated using VCFtools (v0.1.17) [127]. Genome-wide genetic divergence between different Mandarin fish populations was estimated by F_ST_ in overlapping genomic windows (2500 bp, step size: 500 bp) using VCFtools (v0.1.17) [127]. Regions with F_ST_ >= 0.4 and π <= 0.001were identified as divergent genomic regions between *S. chuatsi* and *S. scherzeri*.

### Identification of positively selected genes

Protein sequences of *D. rerio*, *O. niloticus*, *M. zebra*, *L. calcarifer*, *L. crocea*, *D. labrax*, and *S. lucioperca* were downloaded from Ensembl (v104) [97]. Protein sequences of *E. lanceolatus* were downloaded from NCBI. Positively selected genes (PSGs) in the *S. chuatsi* and *S. scherzeri* genomes were identified using PosiGene (v0.1) [128] with parameters “-as = *D.rerio*, -ts = *S.chuatsi* and *S.scherzeri* -rs = *D.rerio*, -nhsbr”. Genes with a *P*-value < 0.05 were identified to have been subject to positive selection. The alignments of PSGs were visualized using Jalview2 [129].

### Structural variation identification

Structural variations (SVs) between the genomes of *S. chuatsi* and *S. scherzeri* were identified using two approaches. First, the SVs were identified by aligning the genome of *S. scherzeri* against the genome of *S. chuatsi* using BLASR [130]. And SVs between the two genomes were identified by combining the results of smartie-sv [56] and SyRI (v4.1) [57]. Second, high-quality Nanopore reads of *S. scherzeri* were aligned against the genome of *S. chuatsi* using NGMLR (v0.2.7) [55]. And Nanopore reads of *S. chuatsi* were aligned against the genome of *S. scherzeri* using NGMLR (v0.2.7). SVs were identified based on both alignment results using Sniffiles (v1.0.11) [55]. SVs with length less than 50 bp were removed. We used Samplot (v1.3.0) to visualize and validate candidate SVs with the alignments of high-quality Nanopore and Illumina reads.

### Histone modification analysis

CUT&Tag assay was performed as described previously with modifications [14]. Briefly, liver cells of *S. chuatsi* and *S. scherzeri* were harvested and gently washed twice in 300 μL wash buffer (20mM HEPES pH 7.5; 150mM NaCl; 0.5mM Spermidine; 1× Protease inhibitor cocktail). A 1:50 dilution of H3K27ac rabbit pAb (ab4729, Abcam), H3K4me3 rabbit pAb (ab8580, Abcam) or H3K4me1 rabbit pAb (ab8895, Abcam) was used as primary antibody for incubation. A 1:50 dilution of Goat Anti-Rabbit IgG (ab8580, Abcam) was used as secondary antibody. For constructing negative control library (IgG), we only added secondary antibody without primary antibody. Cells were washed with Dig-Wash buffer to remove unbound antibodies. A 1:200 dilution of pG-Tn5 adapter complex was added to the cells and incubated with pG-Tn5 protein for 1 hour. Cells were then washed twice in Dig-300 Buffer to remove unbound pG-Tn5 protein. Next, cells were resuspended in tagmentation buffer (10mM MgCl2 in Dig-300 Buffer) and incubated at 37 °C for 1 hour. To stop tagmentation, 10 μL of 0.5M EDTA, 3 μL of 10% SDS and 2.5 μL of 20 mg/mL Proteinase K was added to the sample, which was incubated at 55 °C for 1 hour. DNA was purified using phenol–chloroform–isoamyl alcohol and ethanol, washed with 100% ethanol, and suspended in water. The libraries were amplified by mixing the DNA with 2μL of a universal i5 and uniquely barcoded i7 primer. After DNA quantification and qualification, all libraries were sequenced with 2×150 bp chemistry on the Illumina Nova-seq 6000 platform (Illumina).

CUT&Tag sequencing reads were processed using a pipeline described previously [14]. Low-quality and sequencing-adapter-contaminated Illumina reads were filtered and trimmed with trimmomatic (v0.39) [109]. The filtered reads of *S. chuatsi* and *S. scherzeri* were aligned to the respective genome using Bowtie2 (v2.3.2) [131] with parameters “--local --very-sensitive --no-mixed --no-discordant -I 10 -X 700”. Duplicated reads and reads with mapping quality scores (MAPQ) less than 30 were removed using Samtools (v1.9-52) [112]. MACS2 (v2.1.1) [132] was used to identify both broad and narrow peaks for H3K27ac and H3K4me3 histone modifications with parameters “--keep-dup all”, and only broad peaks for H3K4me1 histone modifications with parameters “--keep-dup all --broad”. Peaks that are significantly differentiated between the genomes of *S. chuatsi* and *S. scherzeri* (|log2FC| >= 1, *P* <= 0.05) were identified using MAnorm (v1.1.4) [133]. Genes that are potentially regulated by the differentiated enhancers and promoters were identified using ChIPseeker [134]. Profiles of CUT&Tag signals of H3K27ac, H3K4me1, and H3K4me3 were visualized with plotProfile function implemented in deepTools (v3.5.1) [135]. Clustering of CUT&Tag signals of three histone marks around the transcription start site (TSS) of genes were visualized with computeMatrix and plotHeatmap functions implemented in deepTools (v3.5.1) [135].

### Open chromatin region identification

Assay for transposase-accessible chromatin using sequencing (ATAC-seq) libraries were constructed using an approach described previously [136]. Liver cells of *S. chuatsi* and *S. scherzeri* were harvested, and the cell suspension was prepared. Cell membranes were lysed to obtain the nucleus. Then, Tn5 transposase was added to cut open DNA. The DNA fragments cut by the enzyme were amplified by PCR and then sequenced with 2X150 bp chemistry on the Illumina HiSeq X Ten platform (Illumina).

Low-quality and sequencing-adapter-contaminated ATAC-seq reads were filtered and trimmed with trimmomatic (v0.39) [109]. The filtered reads of *S. chuatsi* and *S. scherzeri* were aligned to the respective genome using Bowtie2 (v2.3.2) [131] with parameters “--local --very-sensitive --no-mixed --no-discordant -I 10 -X 700”. Duplicated reads and reads with mapping quality scores (MAPQ) less than 30 were removed using Samtools (v1.9-52) [112]. We used MACS2 (v2.1.1) [132] to identify peaks with parameters “--shift -75 --extsize 150 --nomodel -B --SPMR --keep-dup all”.

### Transcriptome sequencing for epigenomic analyses

Samples from liver were collected for *S. chuatsi* and *S. scherzeri* to construct sequencing libraries of RNA-seq for epigenomic analyses. Total RNA was extracted with TRIzol reagent (Invitrogen). Purity and integrity of RNA was monitored by NanoDrop 2000 spectrophotometer (NanoDrop) and Bioanalyzer 2100 system (Agilent). The mRNA was purified from the total RNA using poly-T oligo-attached magnetic beads. Paired-end sequencing libraries were generated from the purified mRNA using the VAHTS Universal V6 RNA-seq Library Kit for MGI (Vazyme) with unique index codes. Sequencing was performed with 2X150 bp chemistry on the MGISEQ 2000 platform (MGI Tech).

Low-quality and sequencing-adapter-contaminated reads of RNA-seq were filtered using SOAPnuke (v2.1.6) [137] with parameters “-n 0.05 -q 0.5 -l 20”. Filtered reads of *S. chuatsi* and *S. scherzeri* were aligned to the respective reference genomes using STAR (v2.7.0) [99]. RSEM (v1.3.3) [138] was used to map and calculate gene expression levels represented as fragments per kilobase of exon per million mapped fragments (FPKM). Differential expression analysis was performed using DESeq2 (R4.1.2) [139].

## Availability of data and materials

Raw reads and genome assemblies are accessible in NCBI under BioProject number PRJNA867131. Assembled genome sequences are accessible under Whole Genome Shotgun project number JAPFEY000000000 (*Siniperca chuatsi*) and JAPFEX000000000 (*Siniperca scherzeri*). Raw reads and genome assemblies are also available at the CNGB Sequence Archive (CNSA) of China National GeneBank DataBase (CNGBdb) with accession number CNP0003680. The genome assembly, related annotation files, and source files for generating figures can be accessed through Figshare at https://doi.org/10.6084/m9.figshare.21385059.

## Abbreviations

CRE: cis-regulatory element
SV: structural variant
ChIP-seq: chromatin immunoprecipitation followed by high throughput sequencing
CUT&Tag: Cleavage Under Targets & Tagmentation
H3K27ac: Histone H3 lysine 27 acetylation
H3K4me3: histone H3 lysine 4 trimethylation
PSG: positively selected gene
BUSCO: Benchmarking Universal Single-Copy Orthologs
TE: transposable element
RNA-seq: RNA sequencing
PSMC: pairwise sequential Markovian coalescent
Ma: million years ago
ka: thousand years ago
MBE: Mid-Brunhes Event
OR: olfactory receptor
CBP/p300: CREB-binding protein and p300
FRiP: fraction of reads in peaks
TSS: transcription start site
OCR: open chromatin regions
KEGG: Kyoto Encyclopedia of Genes and Genomes

## Acknowledgements

We thank Dr. Jinlong Wang for providing resequencing samples of *S. chuatsi*, Dr. Chuanfu Dong and Mr. Fei Li for providing resequencing samples of *S. scherzeri* from Jilin Province, Mr. Gencheng Xian for providing resequencing samples of *S. scherzeri* from Guangdong Province. We gratefully acknowledge the National Supercomputing Center in Guangzhou for provision of computational resources.

## Contributions

M.W. and J.G.H. conceived of the project and designed research; X.Z. and Li Z. sequenced the genomes; G.T., Q.C., and R.J. assembled and annotated the genomes; G.T., Long Z., C.L, and Z.Y. performed the evolutionary analyses; G.T., Y.L., and S.W. performed the epigenomic analyses; M.W., J.G.H., and G.T. wrote the paper with contribution from all authors.

## Ethics declarations

### Ethics approval and consent to participate

All operations of fishes were approved by the Institutional Animal Care and Use Committee of Sun Yat-sen University (Protocol number SYSU-IACUC-2020-B0975). All efforts were made to minimize suffering in animals.

### Consent for publication

Not applicable.

