## Supplementary Materials for "Long-read genome assemblies reveals a *cis*-regulatory landscape associated with phenotypic divergence in two sister *Siniperca* fishes"

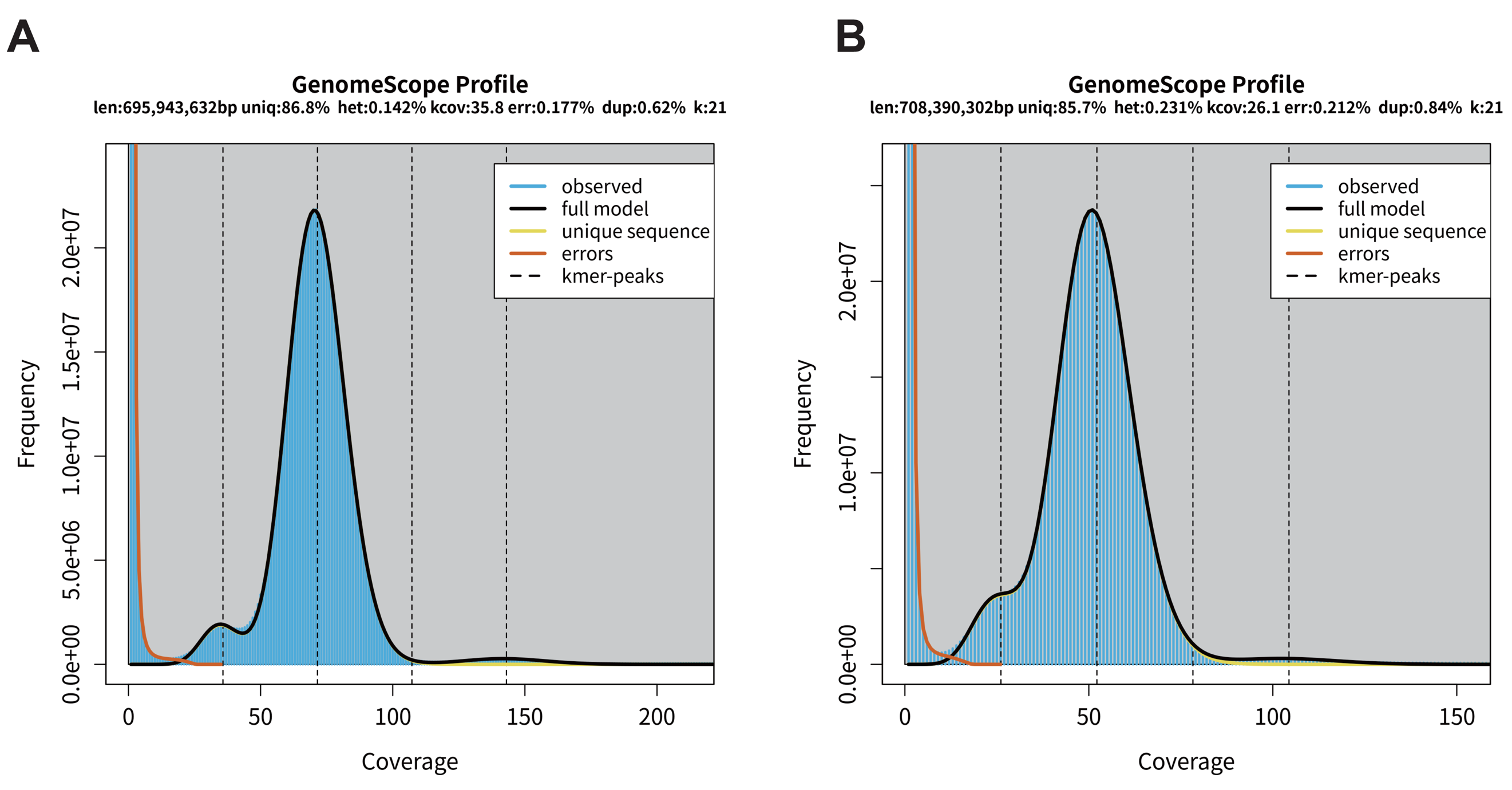

**Supplementary Figure 1. Distribution of 21-mer frequency in *S. chuatsi* (A) and *S. scherzeri* (B) genomes.** The genome sizes of *S. chuatsi* and *S. scherzeri* were estimated to be 695.9 Mb and 708.4 Mb, respectively. The heterozygous rates of *S. chuatsi* genome and *S. scherzeri* genome were estimated to be 0.142% and 0.231%, respectively. (len: haploid length; uniq: percentage non-repetitive sequence; het: percentage estimated heterozygosity; kcov: average *k*-mer coverage for heterozygous bases; err: percentage sequencing error rate; dup: mean read duplication rate; k: *k*-mer size).

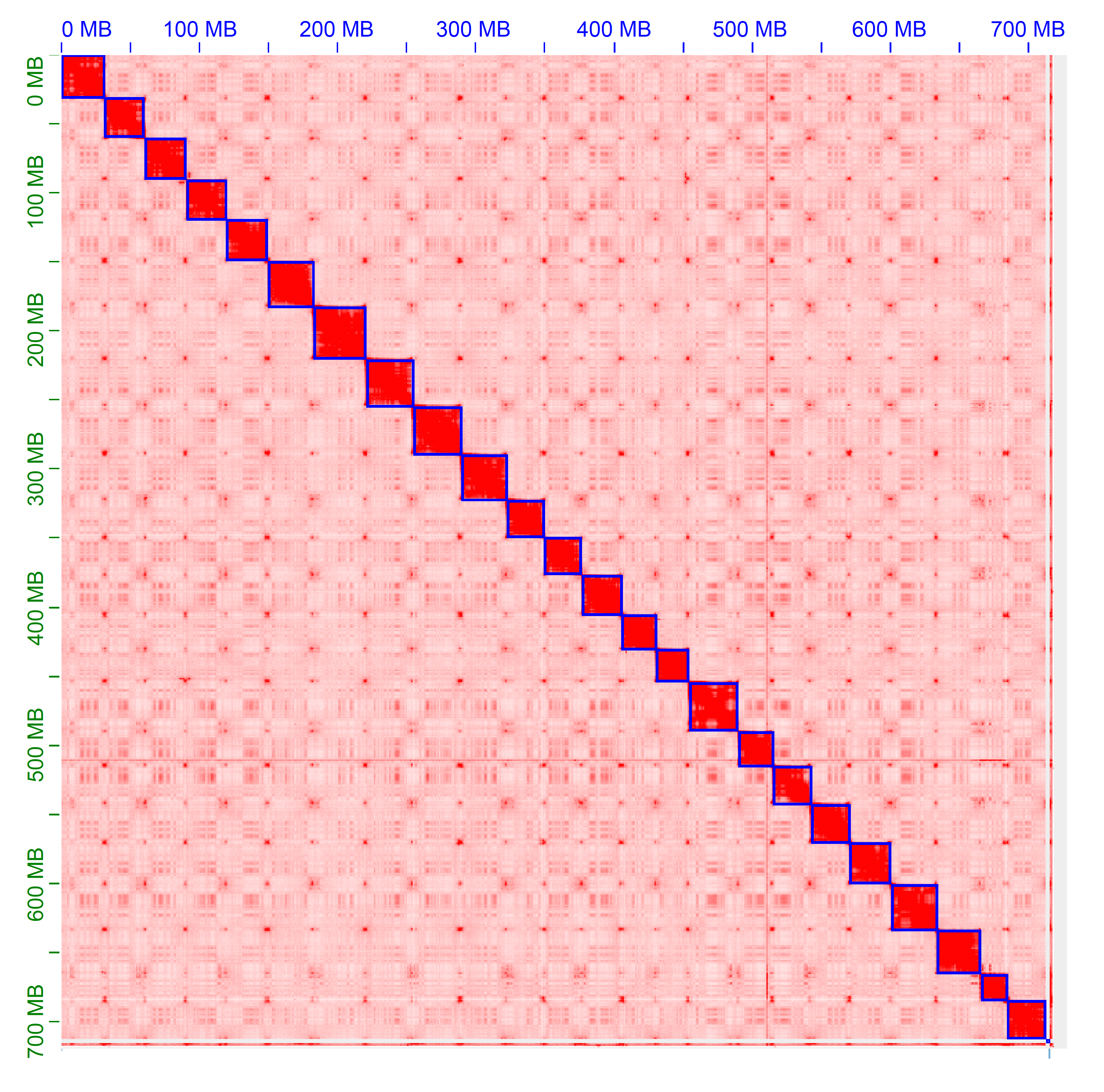

**Supplementary Figure 2 Hi-C interaction matrix for assembled *S. chuatsi* scaffolds generated using the Juicebox Hi-C visualization program.** Darker colors indicate a higher frequency of chromatin interaction. The plot shows clear separation of chromosome boundaries (denoted by blue rectangles) and limited off-diagonal interactions, supporting the global structure of the chromosome-scale scaffolds.

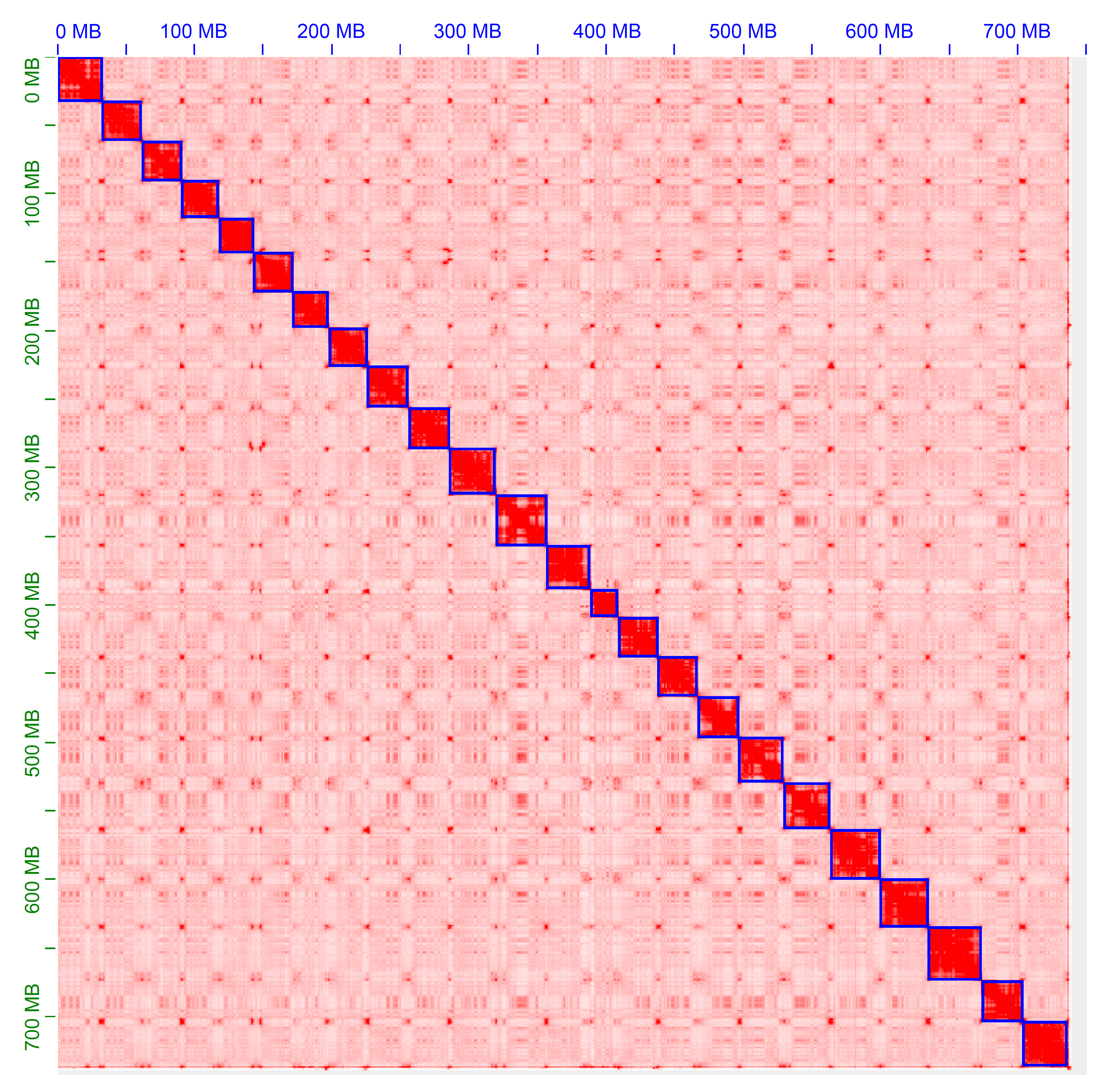

**Supplementary Figure 3 Hi-C interaction matrix for assembled *S. scherzeri* scaffolds generated using the Juicebox Hi-C visualization program.** Darker colors indicate a higher frequency of chromatin interaction. The plot shows clear separation of chromosome boundaries (denoted by blue rectangles) and limited off-diagonal interactions, supporting the global structure of the chromosome-scale scaffolds.

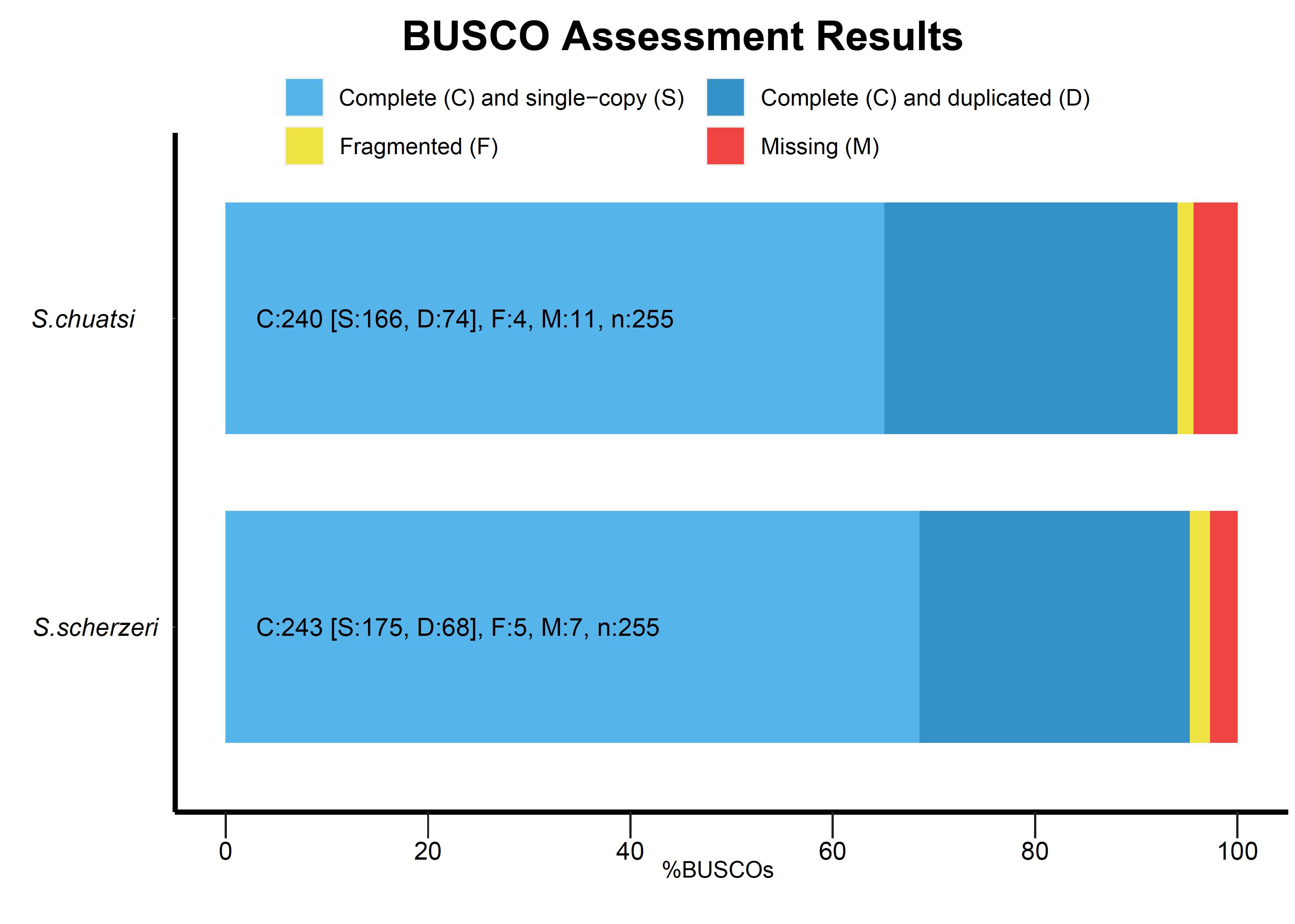

**Supplementary Figure 4. BUSCO assessment of predicted gene model in *S. chuatsi* and *S.scherzeri* .**

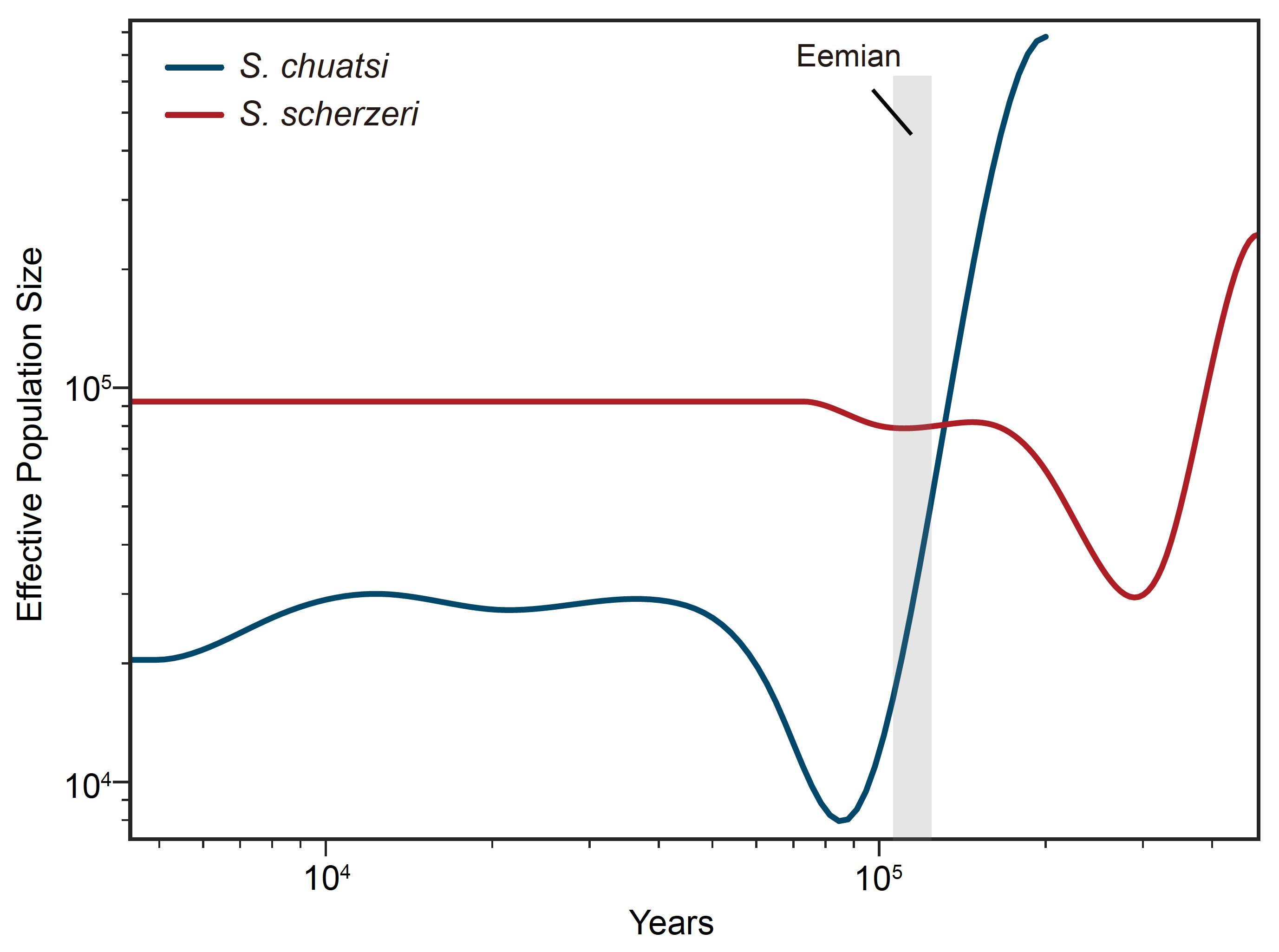

**Supplementary Figure 5 History of ancestral population size of *S. chuatsi* and *S. scherzeri* from north China (Jilin Province).** The ancestral population size of *S. scherzeri* from north China declined at ~300 ka.

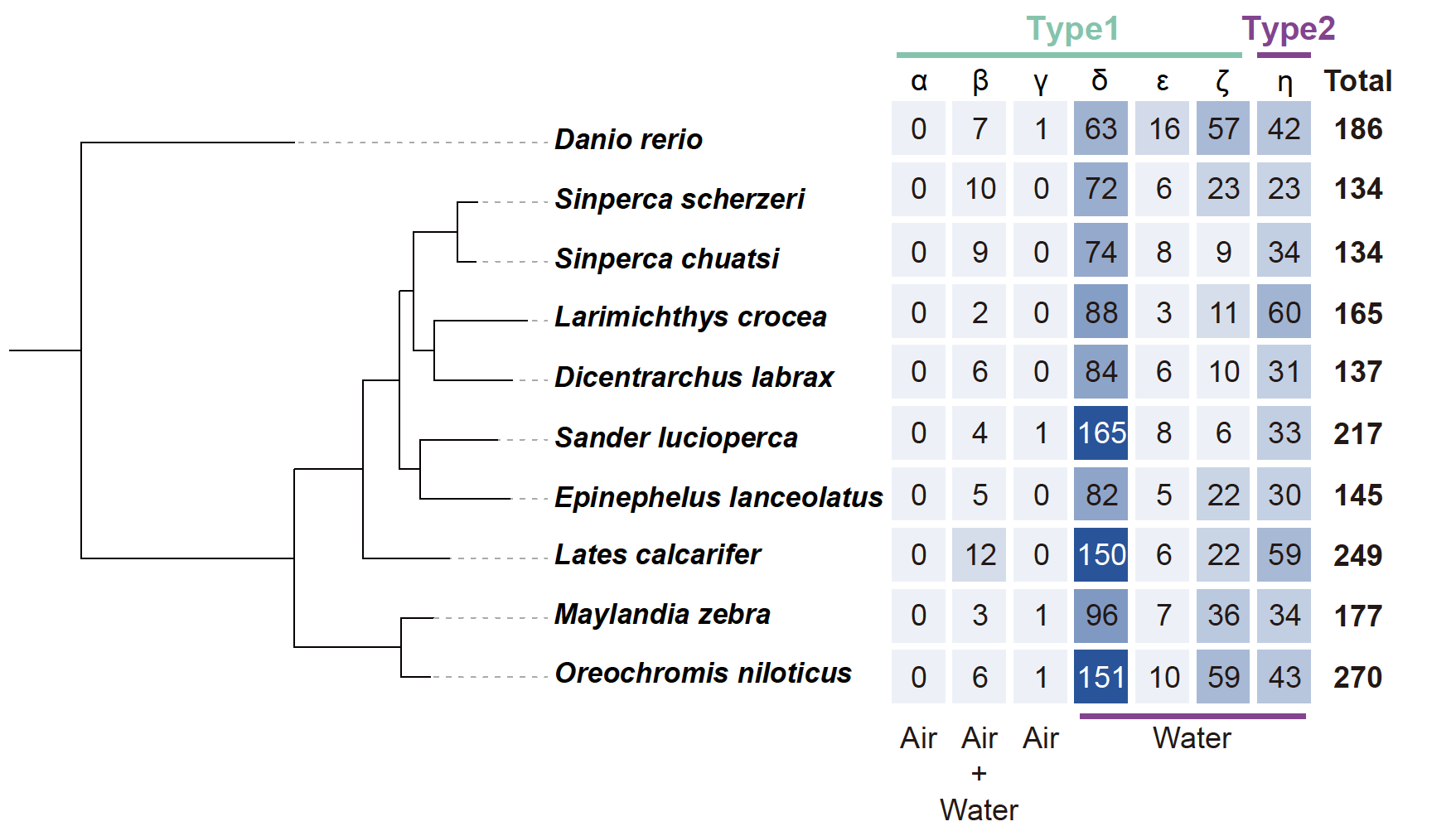

**Supplementary Figure 6 Functional olfactory receptor (OR) genes identified in the genomes of *S. chuatsi* and *S. scherzeri* as well as seven perciform fish species.** *Siniperca chuatsi* and *S. scherzeri* have less OR genes than other perciform fish species.

**OrthoFinder identified ortholog-group of olfactory receptor (OR) genes that are contracted in the genomes of *S. chuatsi* and *S. scherzeri*.** Five ortholog-groups (a, b, c, d, e) belong to three OR subfamilies (*δ*, *ε, ζ*) are contracted in the genomes of *S. chuatsi* and *S. scherzeri*.

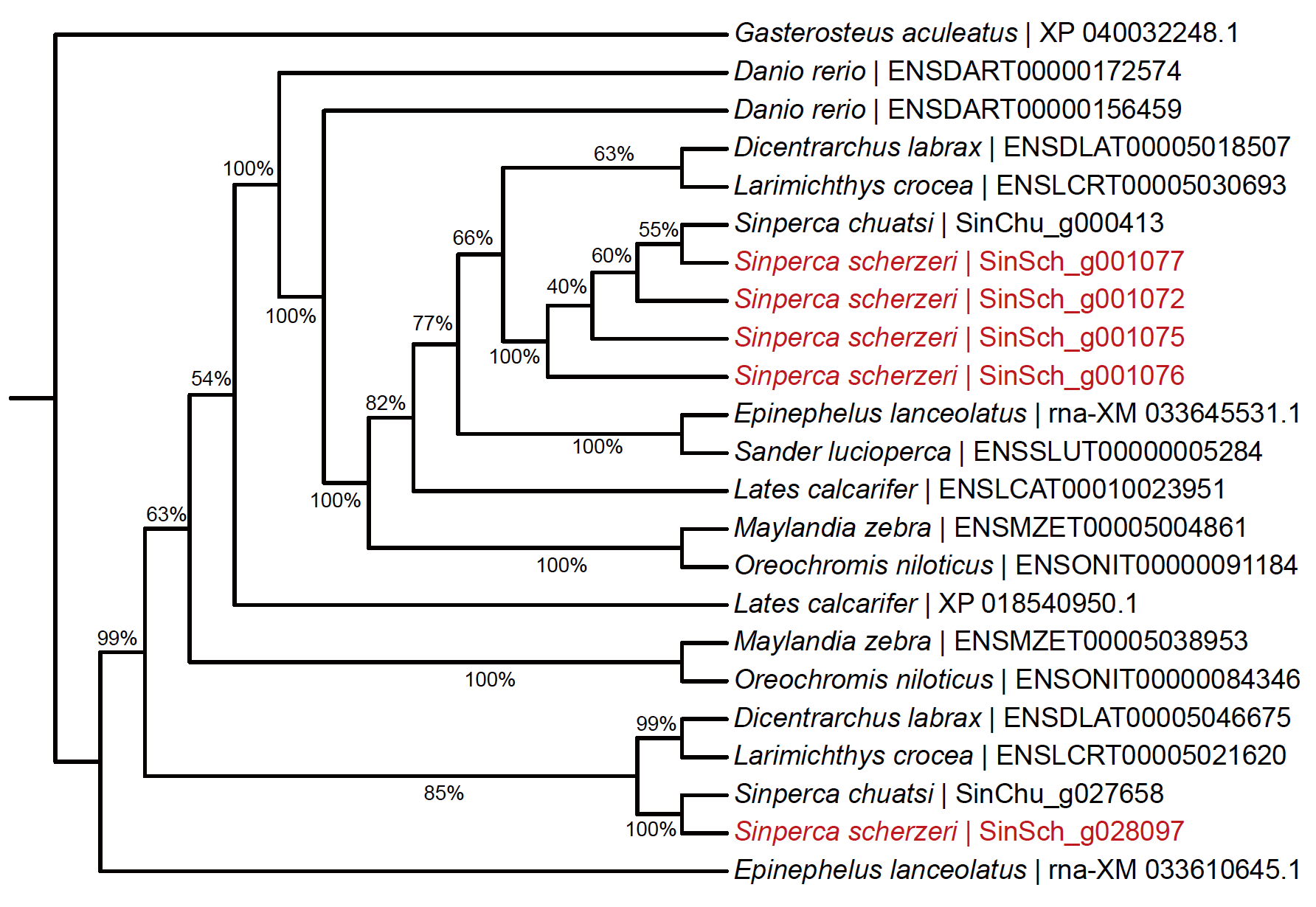

**Supplementary Figure 7 Phylogenetic tree of CBP/p300 genes in ten perciform fishes.** CBP/p300 family was significantly expanded in the genome of *S. scherzeri* (5 copies) compared with all other perciform fishes (2 copies).

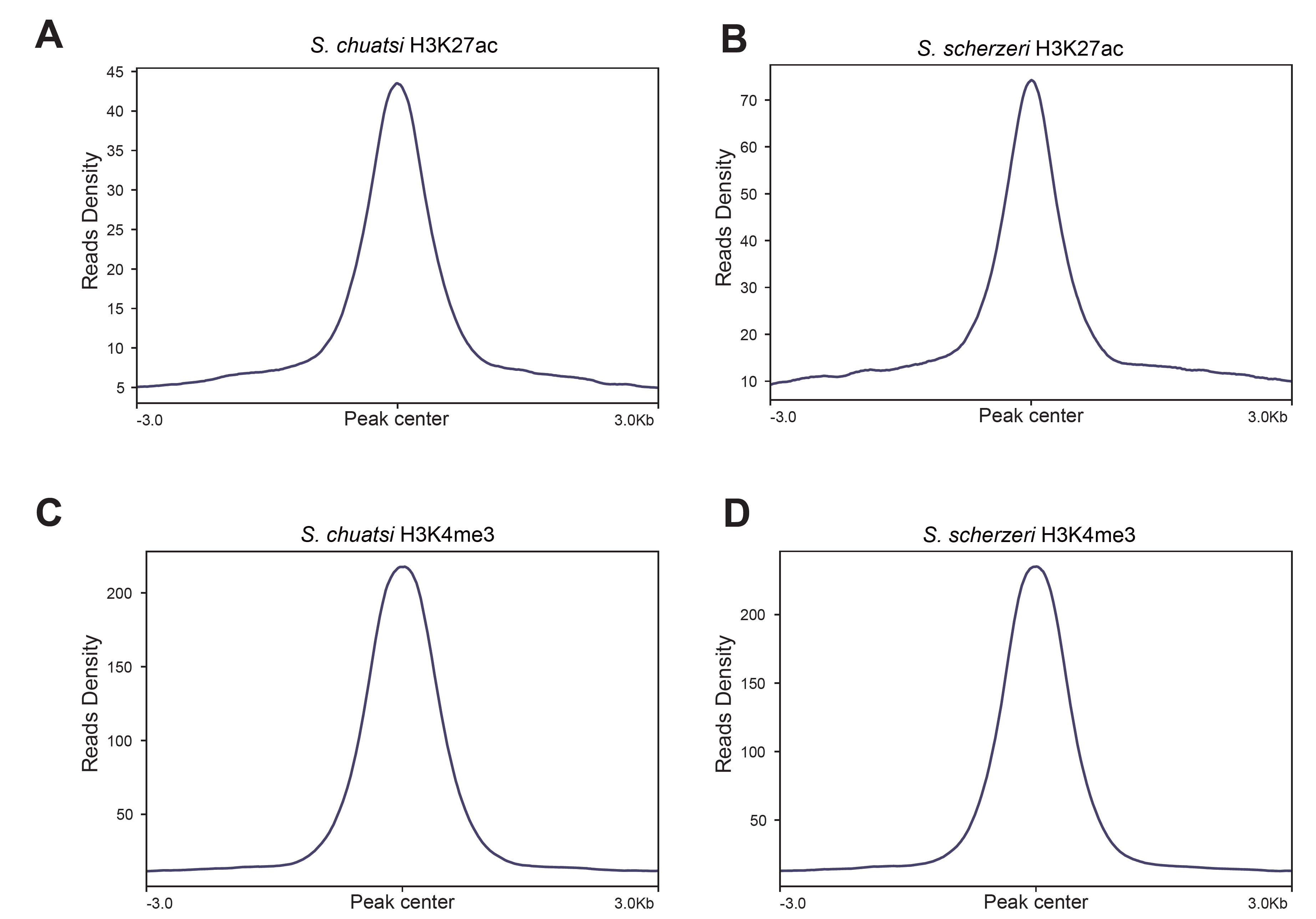

**Supplementary Figure 8 CUT&Tag signals of H3K27ac and H3K4me3.** H3K27ac CUT&Tag signals over replicated H3K27ac peaks in the genomes of *S. chuatsi* (**A**) and *S. scherzeri* (**B**). H3K4me3 CUT&Tag signals over replicated H3K4me3 peaks in the genomes of *S. chuatsi* (**C**) and *S. scherzeri* (**D**).

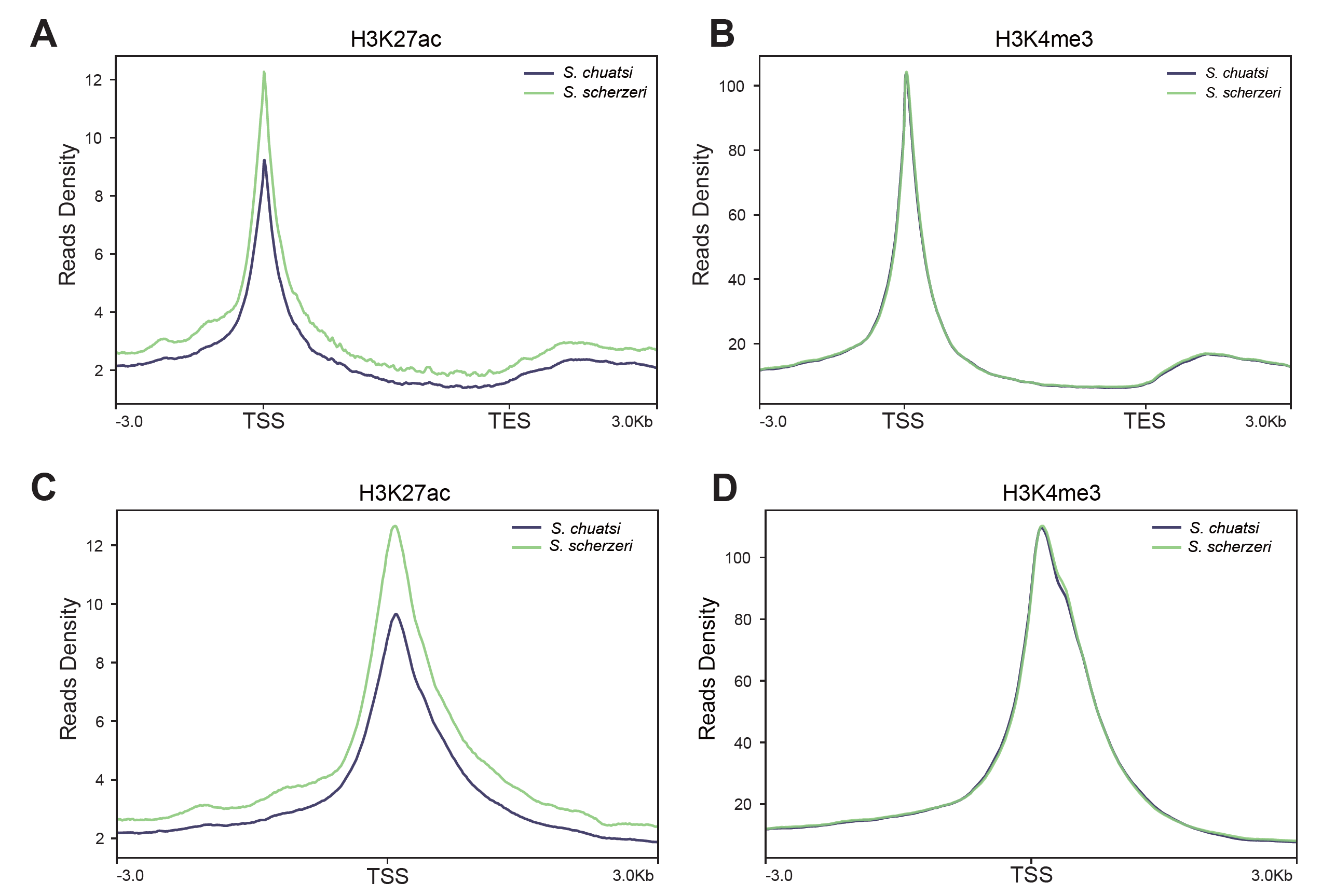

**Supplementary Figure 9. The quality control for CUT&Tag data. A)** The majority of *S. chuatsi* and *S. scherzeri* genes show increased H3K27ac signal. **B)** The majority of *S. chuatsi* and *S. scherzeri* genes show increased H3K4me3 signal. **C)** Transcription start sites (TSS) show particularly high H3K27ac signal in the genomes of *S. chuatsi* and *S. scherzeri*. **D)** Transcription start sites (TSS) show particularly high H3K4me3 signal in the genomes of *S. chuatsi* and *S. scherzeri*.

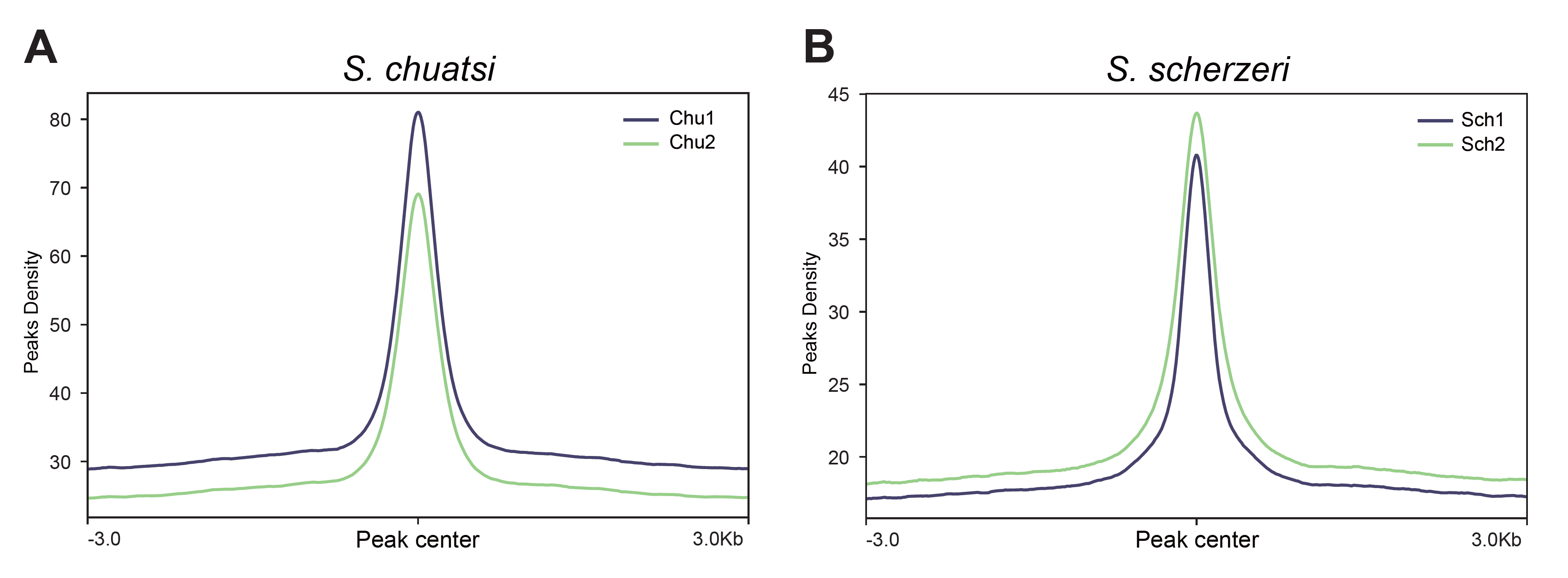

**Supplementary Figure 10 ATAC-seq signals in the genomes of *S. chuatsi* and *S. scherzeri***

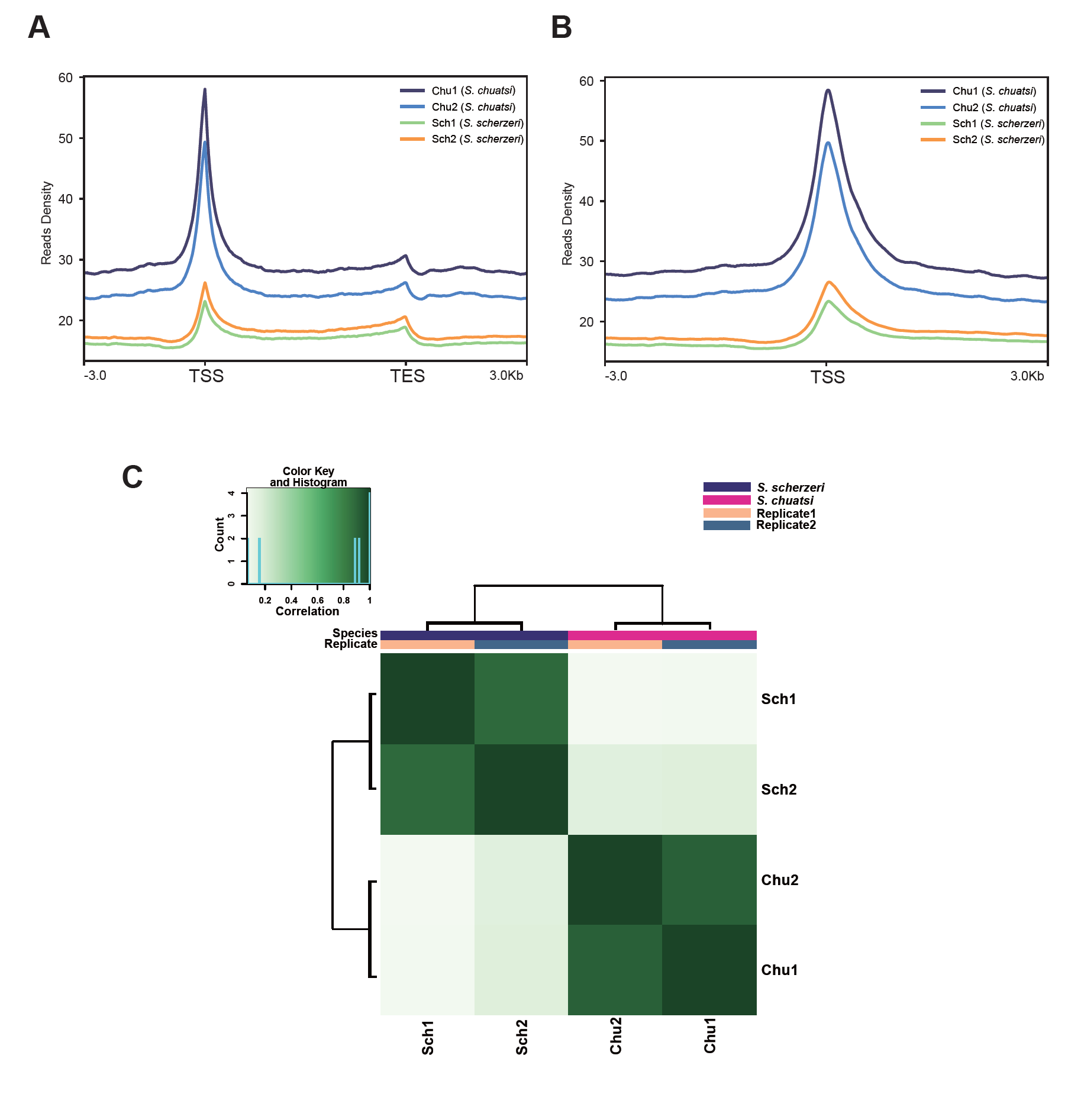

**Supplementary Figure 11 The quality control for ATAC-seq data. A)** The majority of *S. chuatsi* and *S. scherzeri* genes show increased ATAC-seq signal. **B)** Transcription start sites (TSS) show particularly high ATAC-seq signal in the genomes of *S. chuatsi* and *S. scherzeri*. **C)** Correlation between two biological replicates of ATAC-seq of *S. chuatsi* (Chu1, Chu2) and *S. scherzeri* (Sch1, Sch2).

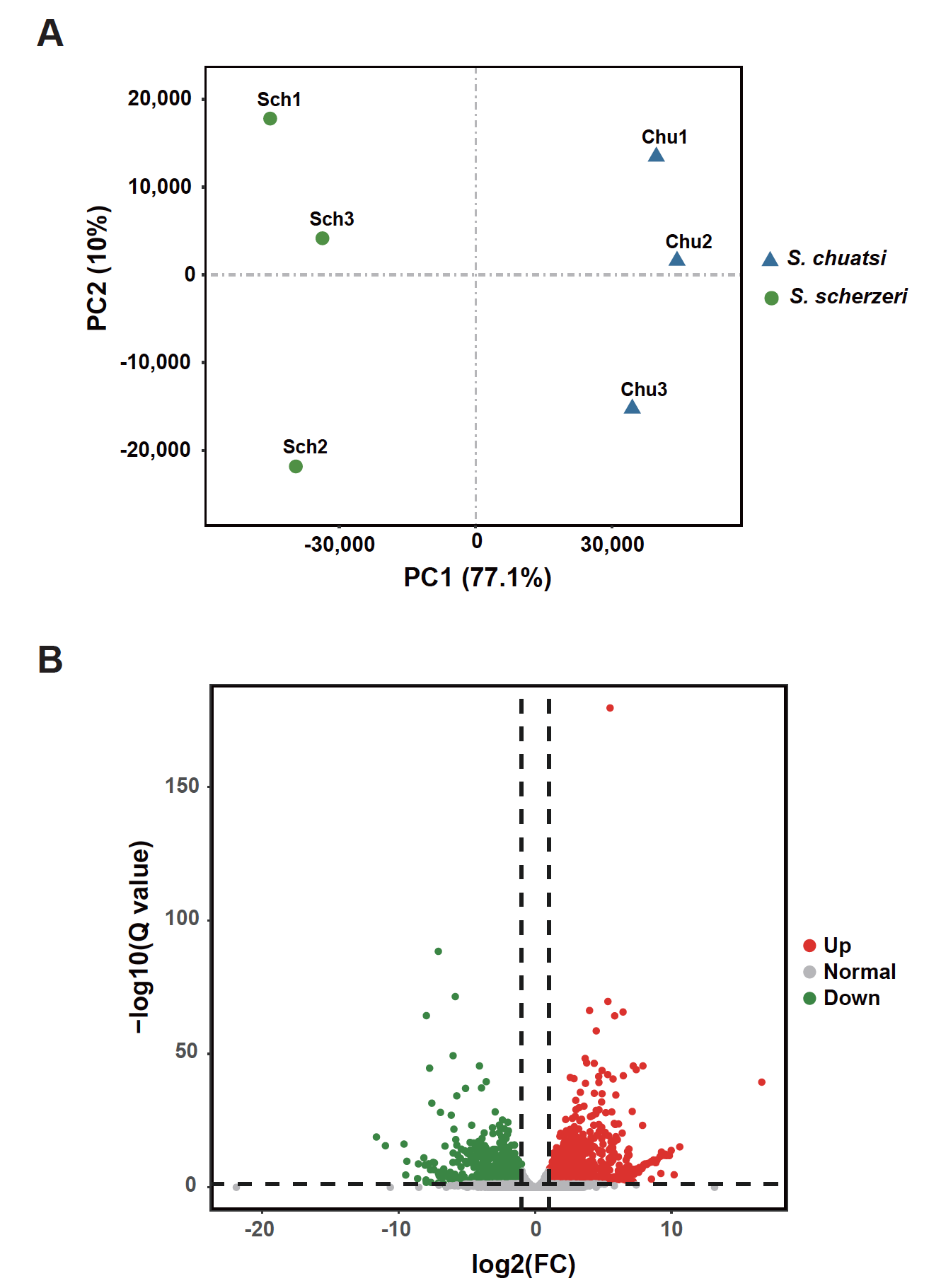

**Supplementary Figure 12 Identification of differentially expressed genes (DEGs) in liver of *S. chuatsi* and *S. scherzeri*. (A)** Principal component analysis (PCA) of RNA-seq data revealed expression divergence between *S. chuatsi* and *S. scherzeri*. **(B)** In total, 2,099 genes are up-regulated, and 1,771 genes are down-regulated in *S. chuatsi* compared with *S. scherzeri*.

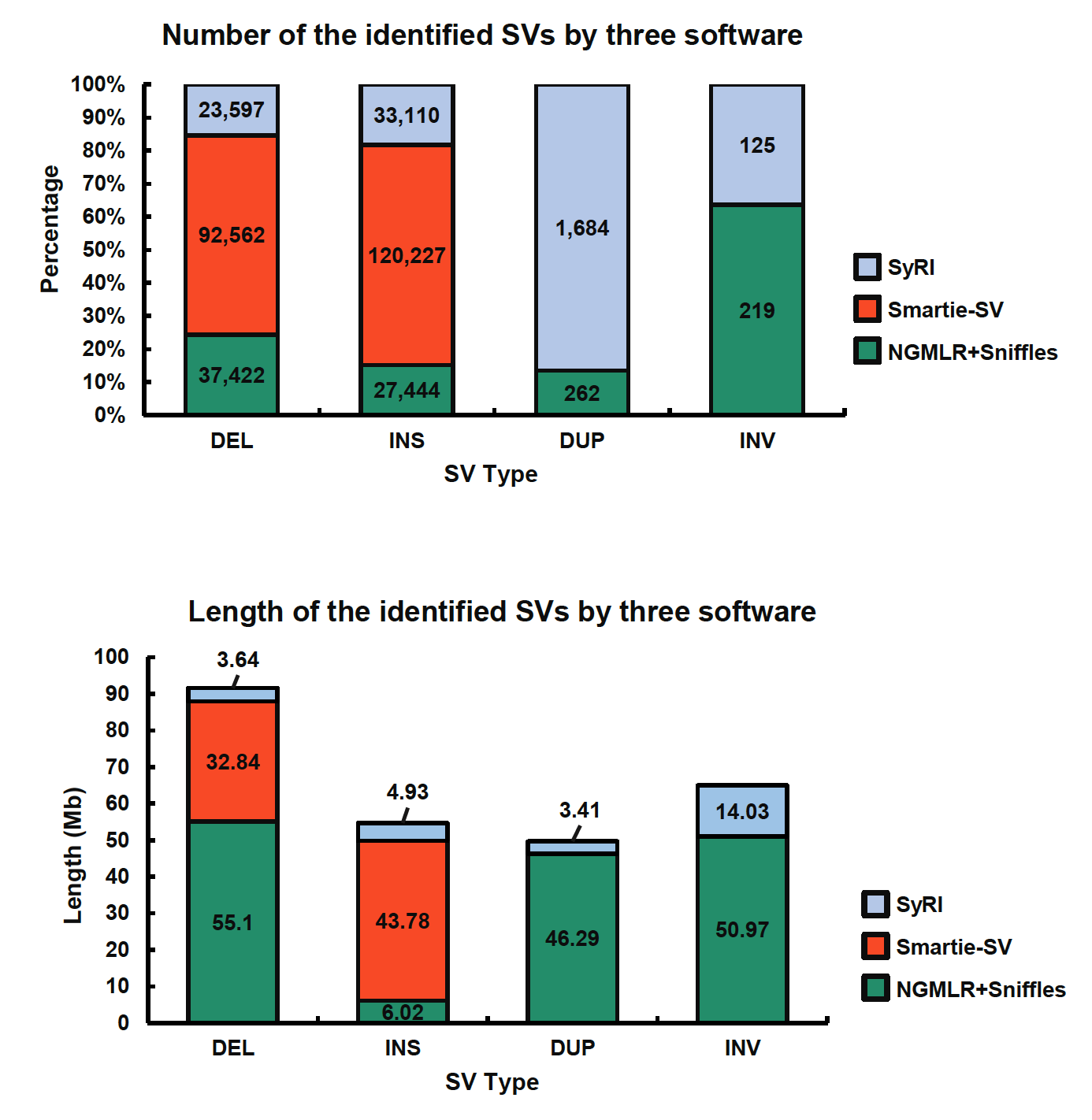

**Supplementary Figure 13 Summary of genomic structural variations (SVs) detection using three different software. A)** Number of identified SVs. **B)** Length of identified SVs.

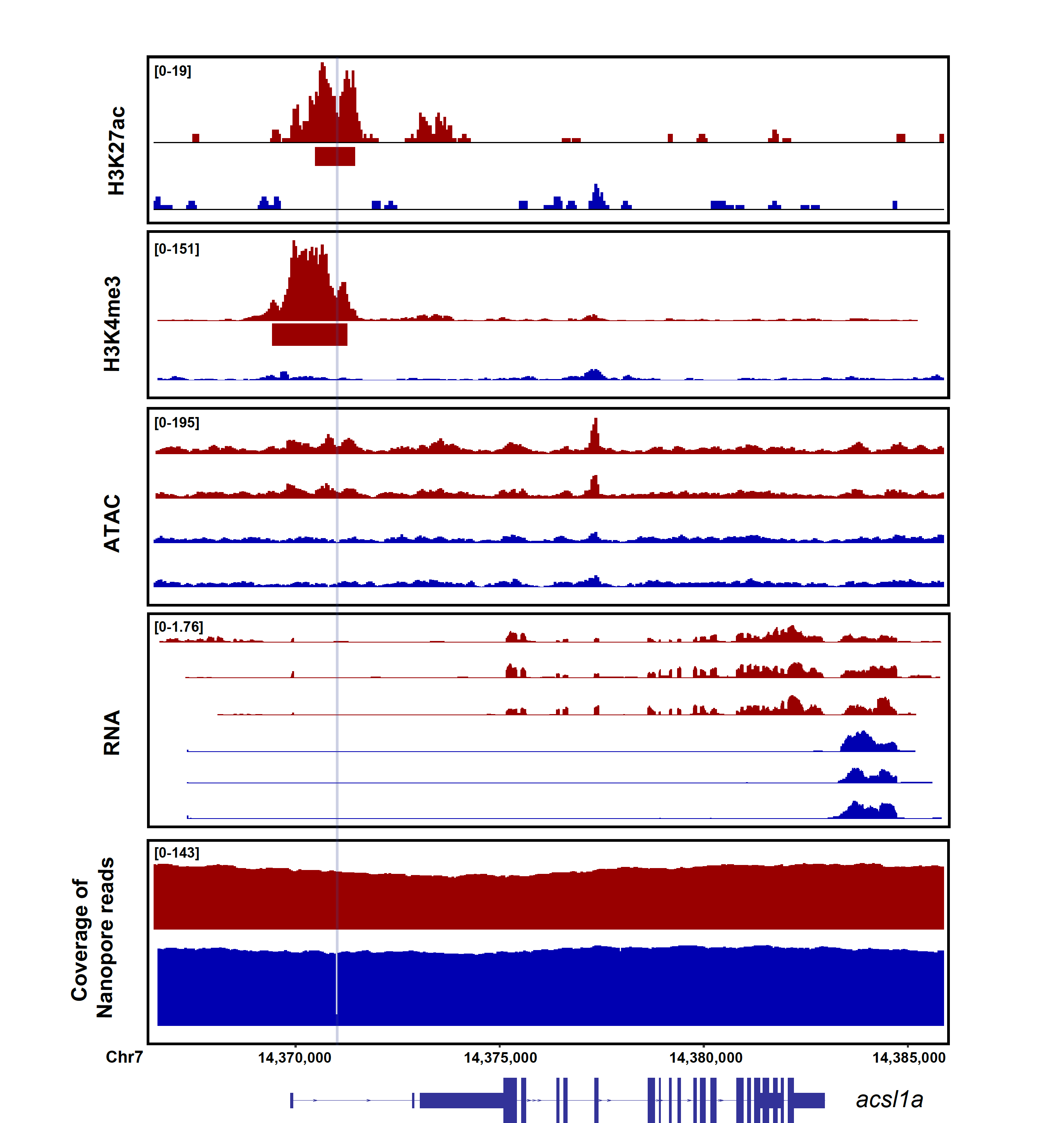

**Supplementary Figure 14 *Cis*-regulatory divergence between *S. chuatsi* (Red) and *S. scherzeri* (Blue) at *acsl1a* gene.** The intensity of H3K27ac, H3K4me3, ATAC-seq, as well as gene expression level are shown. In addition, mapping coverage of Nanopore reads are plotted to indicate genomic structural variants. A deletion in *S. scherzeri* is denoted with a light blue shade. MACS2 identified H3K27ac and H3K4me3 peaks are denoted with rectangles below tracks. Transcripts (with exons as boxes) are depicted.

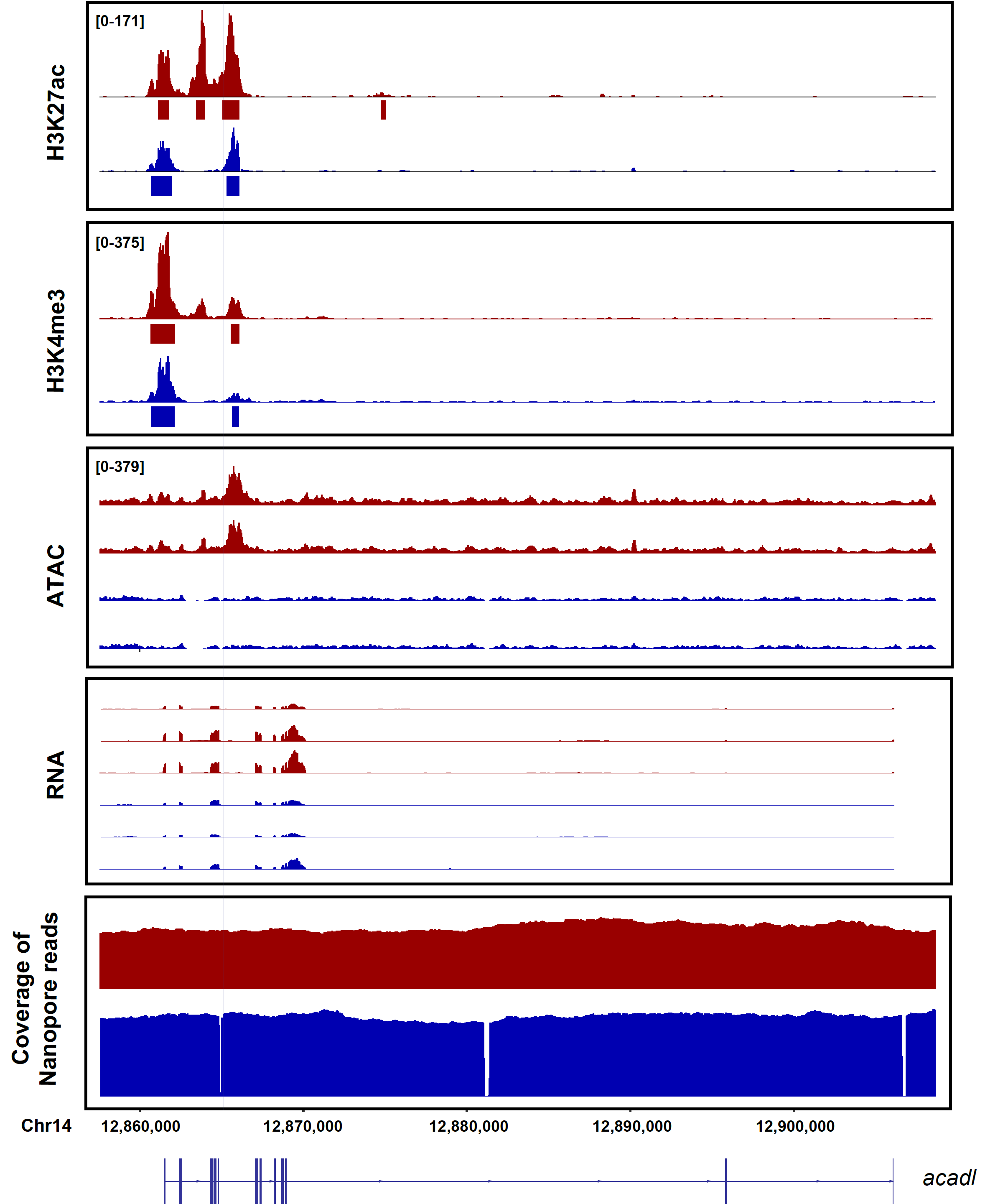

**Supplementary Figure 15 *Cis*-regulatory divergence between *S. chuatsi* (Red) and *S. scherzeri* (Blue) at *acadl* gene.** The intensity of H3K27ac, H3K4me3, ATAC-seq, as well as gene expression level are shown. In addition, mapping coverages of Nanopore reads are plotted to indicate genomic structural variants. A deletion in *S. scherzeri* is denoted with a light blue shade. MACS2 identified H3K27ac and H3K4me3 peaks are denoted with rectangles below tracks. Transcripts (with exons as boxes) are depicted.

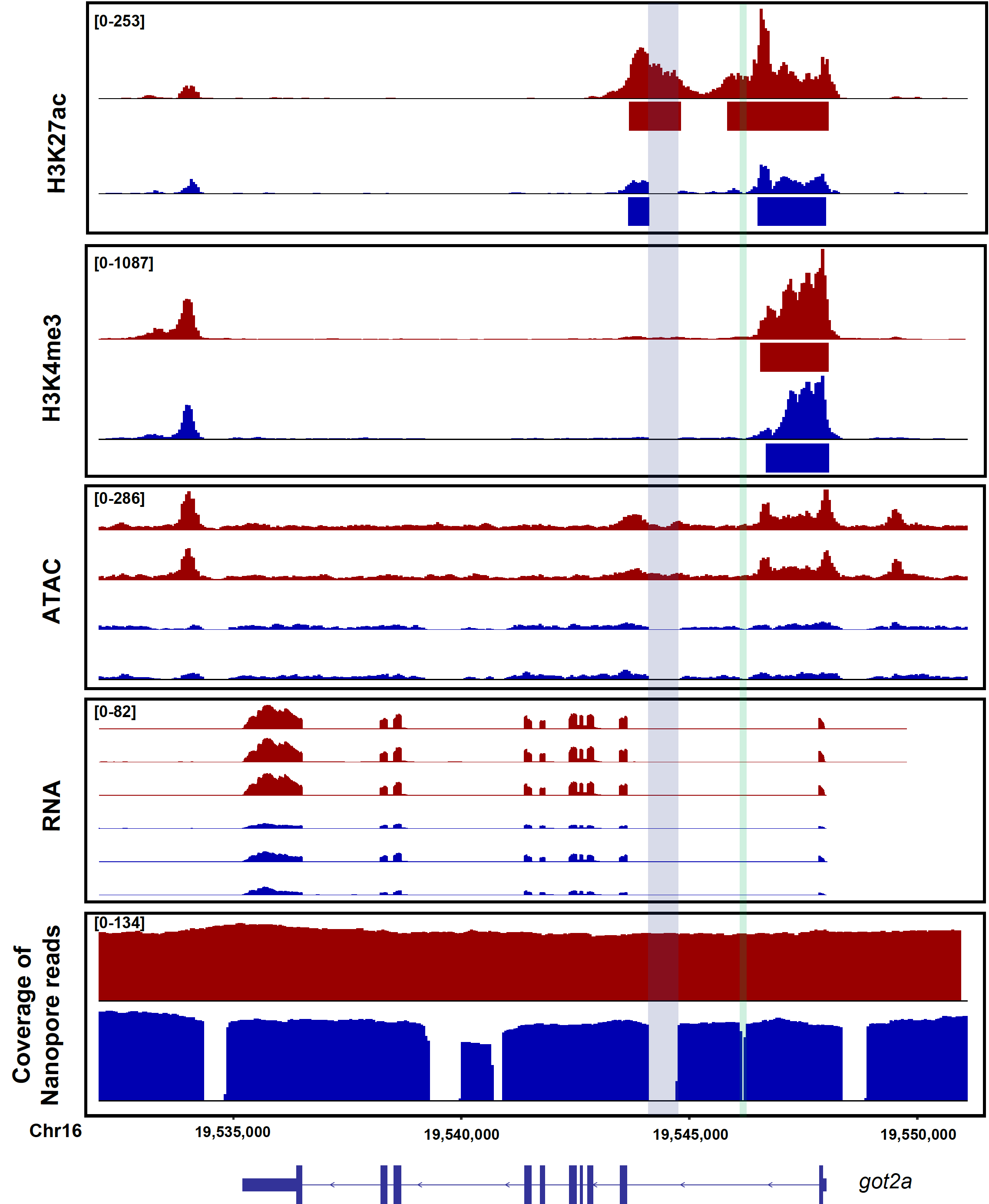

**Supplementary Figure 16 *Cis*-regulatory divergence between *S. chuatsi* (Red) and *S. scherzeri* (Blue) at *got2a* gene.** The intensity of H3K27ac, H3K4me3, ATAC-seq, as well as gene expression level are shown. In addition, mapping coverages of Nanopore reads are plotted to indicate genomic structural variants. Two deletions in *S. scherzeri* are denoted with light blue and green shades. MACS2 identified H3K27ac and H3K4me3 peaks are denoted with rectangles below tracks. Transcripts (with exons as boxes) are depicted.

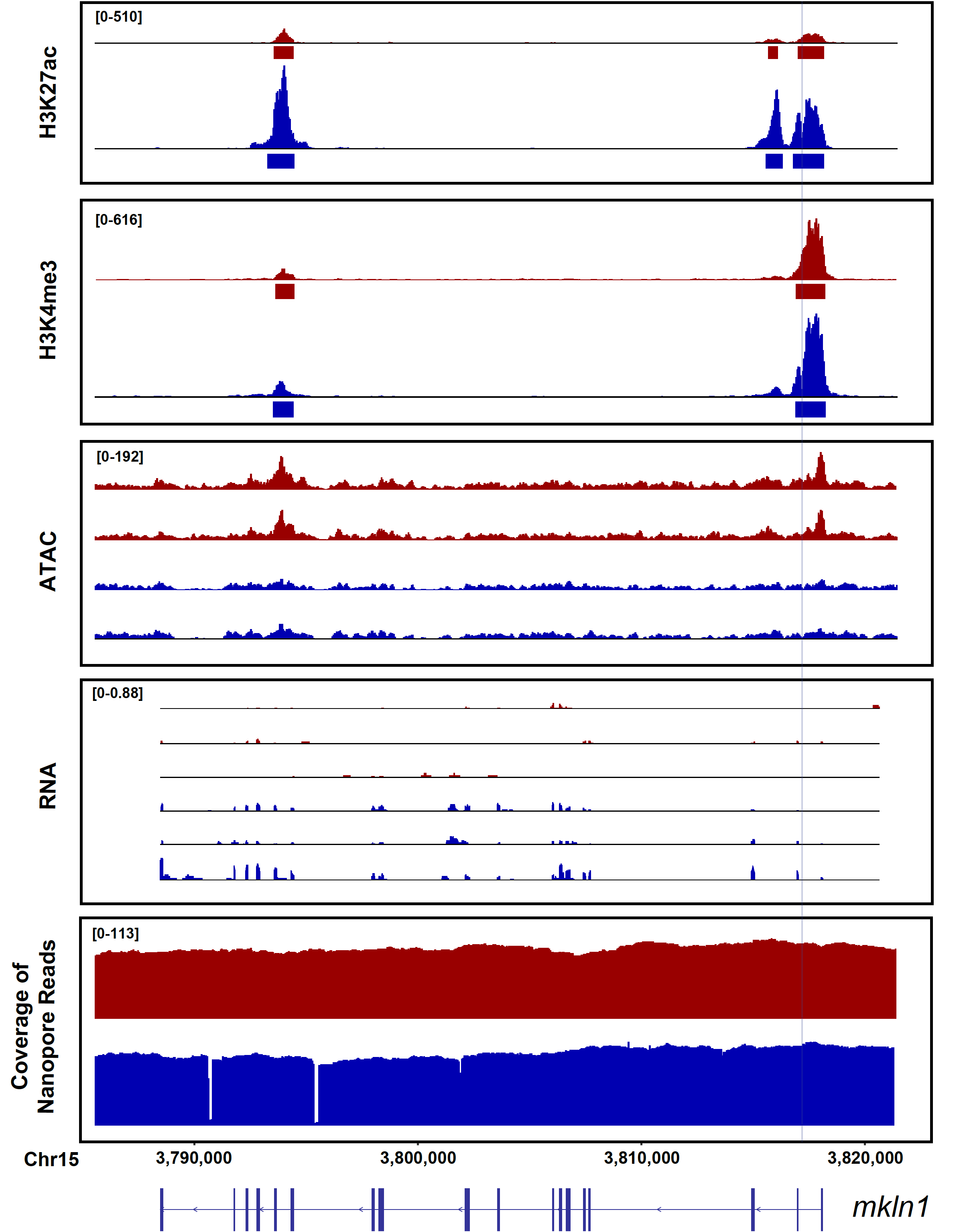

**Supplementary Figure 17 *Cis*-regulatory divergence between *S. chuatsi* (Red) and *S. scherzeri* (Blue) at *mkln1* gene.** The intensity of H3K27ac, H3K4me3, ATAC-seq, as well as gene expression level are shown. In addition, mapping coverages of Nanopore reads are plotted to indicate genomic structural variants. A deletion in *S. chuatsi* is denoted with a light blue shade. MACS2 identified H3K27ac and H3K4me3 peaks are denoted with rectangles below tracks. Transcripts (with exons as boxes) are depicted.

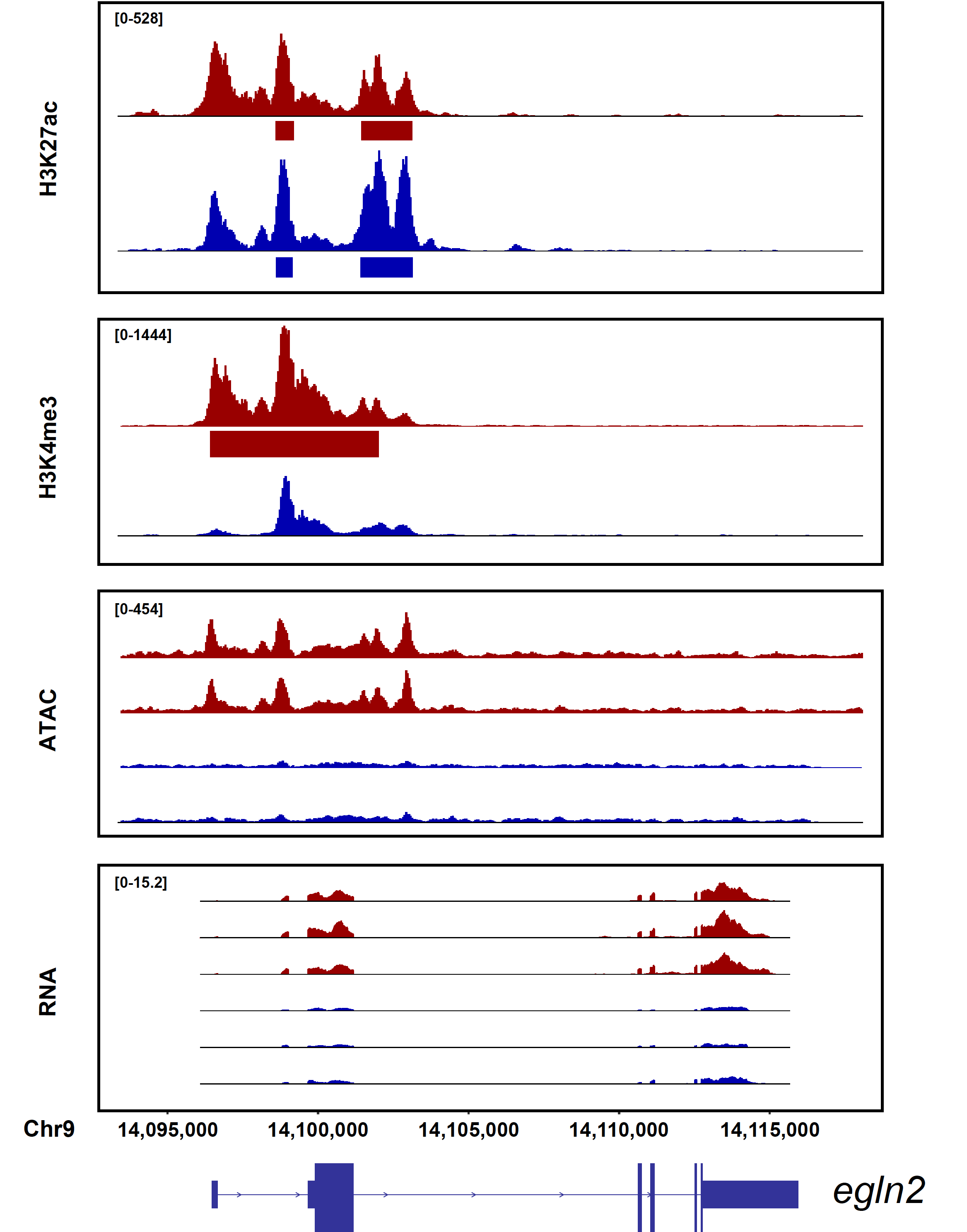

**Supplementary Figure 18 *Cis*-regulatory divergence between *S. chuatsi* (Red) and *S. scherzeri* (Blue) at *egln2* gene.** The intensity of H3K27ac, H3K4me3, ATAC-seq, as well as gene expression level are shown. MACS2 identified H3K27ac and broad H3K4me3 peaks are denoted with rectangles below track. Transcripts (with exons as boxes) are depicted.

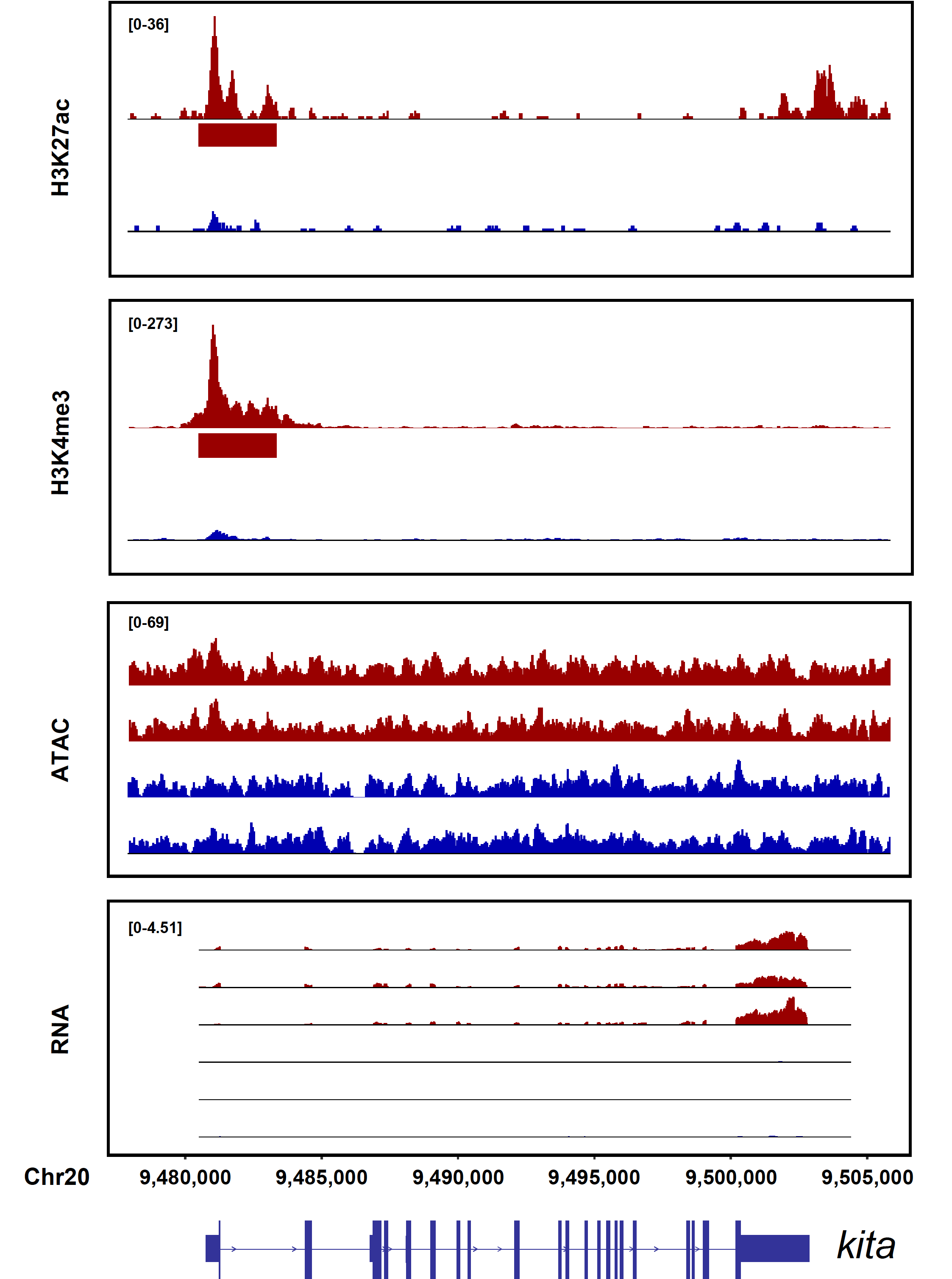

**Supplementary Figure 19 *Cis*-regulatory divergence between *S. chuatsi* (Red) and *S. scherzeri* (Blue) at *kita* gene.** The intensity of H3K27ac, H3K4me3, ATAC-seq, as well as gene expression level are shown. MACS2 identified H3K27ac and broad H3K4me3 peaks are denoted with rectangles below tracks. Transcripts (with exons as boxes) are depicted.

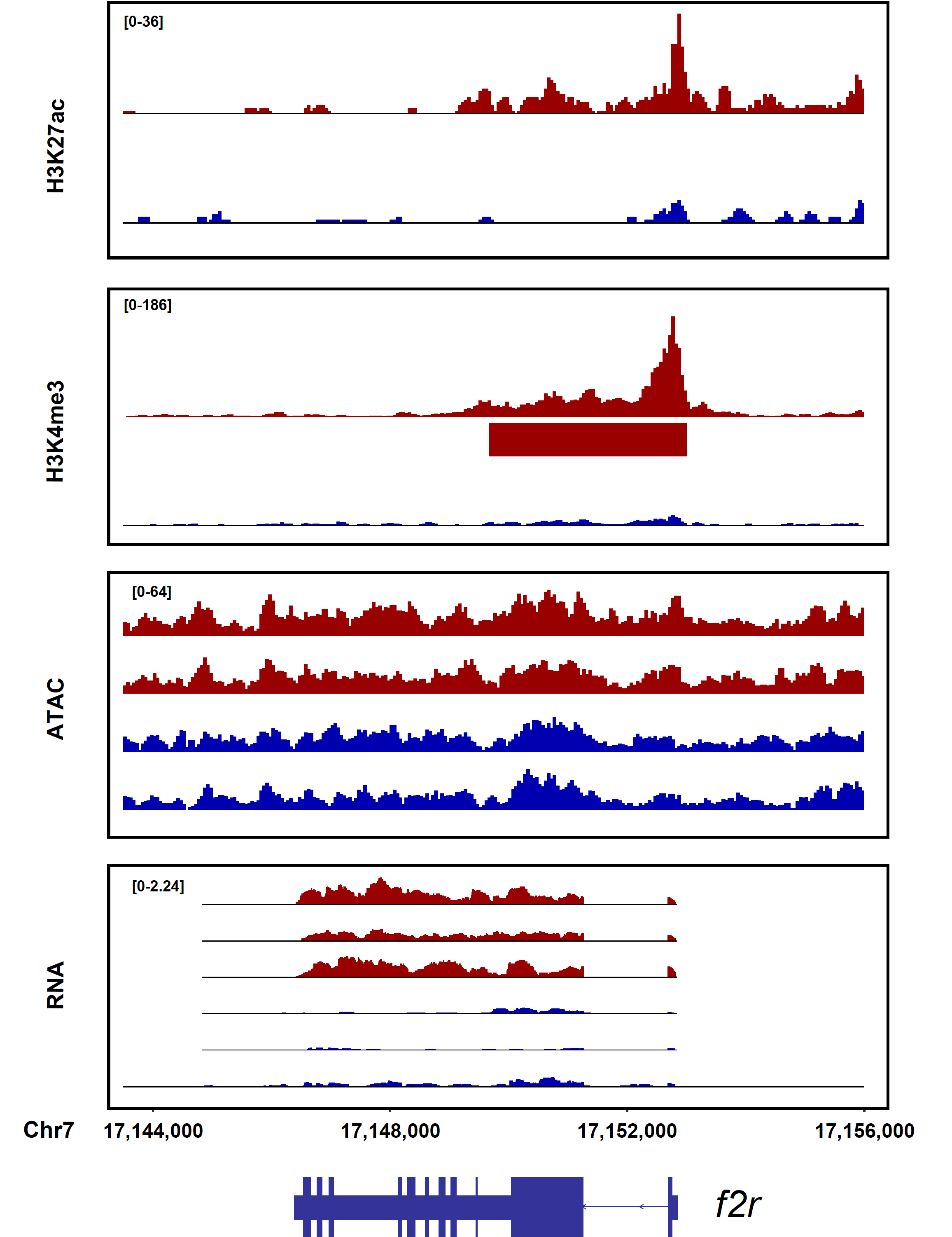

**Supplementary Figure 20 *Cis*-regulatory divergence between *S. chuatsi* (Red) and *S. scherzeri* (Blue) at *f2r* gene.** The intensity of H3K27ac, H3K4me3, ATAC-seq, as well as gene expression level are shown. MACS2 identified H3K27ac, broad H3K4me3 peaks are denoted with rectangles below track. Transcripts (with exons as boxes) are depicted.

**Supplementary Table 1** Basic statistics of Illumina reads

|  | **Number of reads** | **Total bases (bp)** | **Depth of coverage (**$\boldsymbol{\times}$**)** |
| --- | --- | --- | --- |
| *S. chuatsi* | 412,923,280 | 61,938,492,000 | 86.5 |
| *S. scherzeri* | 308,477,640 | 46,271,646,000 | 62.5 |

**Supplementary Table 2** Basic statistics of Nanopore reads

|  | **Total number of reads** | **Total number of bases (Gb)** | **Maximum length of reads (bp)** | **Minimum length of reads (bp)** | **Average length of reads (bp)** | **Depth of coverage (**$\boldsymbol{\times}$**)** |
| --- | --- | --- | --- | --- | --- | --- |
| ***S. chuatsi*** | 4,978,504 | 97.7 | 240,827 | 42 | 19,630 | 136.4 |
| ***S. scherzeri*** | 3,926,889 | 94.3 | 243,874 | 105 | 24,010 | 127.3 |

**Supplementary Table 3** Basic statistics of Hi-C reads

|  | **Total bases (bp)** | **Depth of coverage (**$\boldsymbol{\times}$**)** | **cis-chromosomal interaction**  **(> 20 kb)** | **cis-chromosomal interaction**  **(< 20 kb)** |
| --- | --- | --- | --- | --- |
| *S. chuatsi* | 94,818,391,200 | 132.4 | 24 % | 7 % |
| *S. scherzeri* | 102,637,460,100 | 138.6 | 19 % | 7% |

**Supplementary Table 4** Statistics of *S. chuatsi* genome assembly

|  | Contig | | Scaffold | |
| --- | --- | --- | --- | --- |
|  | Size (bp) | Number | Size (bp) | Number |
| N50 | 21,551,097 | 14 | 29,956,575 | 11 |
| N60 | 19,623,084 | 18 | 29,175,850 | 14 |
| N70 | 13,306,200 | 22 | 28,362,969 | 16 |
| N80 | 8,640,736 | 29 | 27,500,176 | 19 |
| N90 | 3,004,397 | 42 | 24,860,938 | 21 |
| Total | 716,210,655 | 328 | 716,348,155 | 191 |

**Supplementary Table 5** Statistics of *S. scherzeri* genome assembly

|  | Contig | | Scaffold | |
| --- | --- | --- | --- | --- |
|  | Size (bp) | Number | Size (bp) | Number |
| N50 | 16,036,871 | 15 | 30,486,584 | 11 |
| N60 | 14,480,942 | 19 | 30,021,833 | 14 |
| N70 | 9,520,673 | 26 | 29,387,193 | 16 |
| N80 | 5,357,922 | 36 | 28,699,791 | 19 |
| N90 | 3,383,819 | 53 | 27,297,733 | 21 |
| Total | 740,384,376 | 406 | 740,538,376 | 252 |

**Supplementary Table 6** BUSCO evaluation of *S. chuatsi* and *S. scherzeri* genome assemblies

|  | *S. chuatsi* | *S. scherzeri* |
| --- | --- | --- |
| Complete BUSCOs | 3562 (97.9%) | 3593 (98.7%) |
| Complete and single-copy BUSCOs | 3540 (97.3%) | 3567 (98.0%) |
| Complete and duplicated BUSCOs | 22 (0.6%) | 26 (0.7%) |
| Fragmented BUSCOs | 16 | 17 |
| Missing BUSCOs | 62 | 30 |
| Total BUSCO groups searched | 3640 | 3640 |

**Supplementary Table 7** RNA-seq reads alignment statistics of two *S. chuatsi* assemblies

|  | **Assembly** | **Unmapped reads (%)** | **Uniquely mapped reads (%)** | **Multi-mapping reads (%)** | **Mapping rate (%)** |
| --- | --- | --- | --- | --- | --- |
| **Eye** | sinChu7 | 15.50% | 81.50% | 2.42% | 86.13% |
|  | This assembly | 6.79% | 86.72% | 5.88% | 95.00% |
| **Gill** | sinChu7 | 20.19% | 75.80% | 3.48% | 81.64% |
|  | This assembly | 7.23% | 85.40% | 6.86% | 95.10% |
| **Heart** | sinChu7 | 14.38% | 82.07% | 3.17% | 87.52% |
|  | This assembly | 6.85% | 87.71% | 5.05% | 95.17% |
| **Intestine** | sinChu7 | 20.03% | 76.12% | 3.42% | 82.23% |
|  | This assembly | 9.05% | 84.01% | 6.56% | 93.71% |
| **Kidney** | sinChu7 | 14.61% | 82.01% | 2.87% | 87.22% |
|  | This assembly | 9.09% | 86.13% | 4.28% | 93.21% |
| **Liver** | sinChu7 | 6.56% | 86.74% | 6.38% | 94.57% |
|  | This assembly | 5.59% | 88.67% | 5.40% | 95.80% |
| **Spleen** | sinChu7 | 7.94% | 84.68% | 6.96% | 94.42% |
|  | This assembly | 7.39% | 85.05% | 7.15% | 95.36% |
| **Stomach** | sinChu7 | 19.85% | 75.06% | 4.78% | 81.72% |
|  | This assembly | 7.51% | 83.84% | 8.36% | 94.51% |
| **Testis** | sinChu7 | 9.31% | 86.75% | 3.43% | 92.57% |
|  | This assembly | 6.73% | 88.36% | 4.35% | 95.39% |
| **Brain***  **(SRR10867828)** | sinChu7 | 28.86% | 67.91% | 1.74% | 78.14% |
|  | This assembly | 21.72% | 72.20% | 4.52% | 86.40% |
| **Intestine***  **(SRR10867827)** | sinChu7 | 36.82% | 59.18% | 2.68% | 69.86% |
|  | This assembly | 22.28% | 67.67% | 8.65% | 86.76% |
| **Liver***  **(SRR10867826)** | sinChu7 | 33.45% | 59.68% | 5.52% | 72.34% |
|  | This assembly | 19.70% | 67.72% | 11.12% | 88.45% |
| **Muscle***  **(SRR10867825)** | sinChu7 | 31.39% | 58.95% | 8.38% | 75.42% |
|  | This assembly | 25.37% | 65.10% | 8.16% | 84.25% |

**Note:** RNA-seq reads from He *et al.* (2019) [1] were denoted with *

**Supplementary Table 8** RNA-seq reads alignment statistics of two *S. scherzeri* assemblies

|  | Assembly | Unmapped reads (%) | Uniquely mapped reads (%) | Multi-mapping reads (%) | Mapping rate (%) |
| --- | --- | --- | --- | --- | --- |
| Eye | sinSch6b | 6.17% | 89.64% | 3.91% | 96.09% |
|  | This assembly | 5.39% | 91.81% | 2.52% | 96.74% |
| Gill | sinSch6b | 10.35% | 86.09% | 3.20% | 92.71% |
|  | This assembly | 7.16% | 88.14% | 4.35% | 95.44% |
| Heart | sinSch6b | 6.77% | 87.17% | 5.77% | 95.61% |
|  | This assembly | 5.46% | 90.75% | 3.48% | 96.74% |
| Intestine | sinSch6b | 10.29% | 85.27% | 4.03% | 93.01% |
|  | This assembly | 6.92% | 88.92% | 3.79% | 95.83% |
| Kidney | sinSch6b | 8.65% | 85.76% | 5.28% | 94.18% |
|  | This assembly | 6.53% | 89.94% | 3.20% | 96.02% |
| Liver | sinSch6b | 8.16% | 85.66% | 5.75% | 94.26% |
|  | This assembly | 4.55% | 88.77% | 6.39% | 97.29% |
| Spleen | sinSch6b | 11.06% | 85.30% | 3.27% | 92.60% |
|  | This assembly | 8.10% | 87.76% | 3.78% | 95.10% |
| Stomach | sinSch6b | 7.05% | 88.67% | 3.96% | 95.32% |
|  | This assembly | 4.98% | 90.95% | 3.76% | 97.01% |
| Testis | sinSch6b | 7.91% | 86.49% | 5.35% | 94.78% |
|  | This assembly | 5.94% | 90.74% | 3.07% | 96.40% |

**Supplementary Table 9** Statistics of assembly quality and completeness evaluation using Mercury

|  |  | **QV** | **Error rate** | **Completeness (%)** |
| --- | --- | --- | --- | --- |
| *S. chuatsi* | This assembly | 36.9142 | 0.000203508 | 96.397 |
|  | sinChu7 | 32.3679 | 0.000579707 | 97.1919 |
| *S. scherzeri* | This assembly | 35.7313 | 0.000267218 | 95.7469 |
|  | sinSch6 | 30.9618 | 0.000801352 | 93.7093 |

**Supplementary Table 10** Summary of annotated repeats in *S. chuatsi* genome

|  | Number | Length (bp) | Percentage (%) |
| --- | --- | --- | --- |
| **Retroelements** |  |  |  |
| SINEs: | 25,488 | 2,507,374 | 0.35 |
| MIRs | 8,935 | 1,040,905 | 0.15 |
| LINEs: | 151,607 | 37,269,631 | 5.20 |
| LINE1 | 2,091 | 351,931 | 0.05 |
| LINE2 | 73,943 | 17,149,829 | 2.39 |
| LTR elements: | 24,221 | 6,034,393 | 0.84 |
| ERV_classI | 1,217 | 622,070 | 0.09 |
| **DNA transposons：** | **383,869** | **55,184,211** | **7.70** |
| hAT-Charlie | 18,456 | 2,008,459 | 0.28 |
| **Unclassified:** | **600,490** | **76,225,084** | **10.64** |
| **Total interspersed repeats** |  | **177,220,693** | **24.74** |
| **Small RNA:** | **4,196** | **441,630** | **0.06** |
| **Satellites:** | **643** | **169,440** | **0.02** |
| **Simple repeats:** | **318,886** | **16,965,675** | **2.37** |
| **Low complexity** | **37,739** | **2,290,874** | **0.32** |
| **Total** |  | **196,347,086** | **27.41** |

**Supplementary Table 11** Summary of annotated repeats in *S. scherzeri* genome

|  | Number | Length (bp) | Percentage (%) |
| --- | --- | --- | --- |
| **Retroelements** |  |  |  |
| SINEs: | 28,055 | 2,578,008 | 0.35 |
| MIRs | 11,449 | 1,294,403 | 0.17 |
| LINEs: | 154,990 | 43,019,685 | 5.81 |
| LINE1 | 2382 | 514,798 | 0.07 |
| LINE2 | 77,893 | 19,455,186 | 2.63 |
| LTR elements: | 34,632 | 7,746,310 | 1.05 |
| ERV_classI | 4128 | 1,000,999 | 0.14 |
| ERV_classII | 570 | 86,334 | 0.01 |
| **DNA transposons：** | **393,505** | **55,646,827** | **7.51** |
| hAT-Charlie | 12,997 | 1,541,780 | 0.21 |
| TcMar-Tigger | 957 | 69,671 | 0.01 |
| **Unclassified:** | **645,653** | **80,168,797** | **10.83** |
| **Total interspersed repeats** |  | **189,159,627** | **25.54** |
| **Small RNA:** | **3604** | **646,701** | **0.09** |
| **Satellites:** | **2528** | **273,497** | **0.04** |
| **Simple repeats:** | **329,210** | **17,681,281** | **2.39** |
| **Low complexity** | **38,168** | **2,295,851** | **0.31** |
| **Total** |  | **209,398,734** | **28.28** |

**Supplementary Table 12** Whole-genome gene annotation for *S. chuatsi* using different prediction approaches

| **Gene set** | | **Gene Number** | **Exon Number** |
| --- | --- | --- | --- |
| *De novo* based | Braker2 (Illumina) | 38,330 | 392,512 |
|  | Braker2 (PacBio Iso-seq) | 37,689 | 246,953 |
| Homolog based | *Danio rerio* | 31,454 | 290,167 |
|  | *Oryzias latipes* | 39,946 | 353,988 |
|  | *Takifugu rubripes* | 25,652 | 252,838 |
|  | *Gasterosteus aculeatus* | 32,183 | 280,964 |
|  | *Tetraodon nigroviridi* | 21,112 | 229,856 |
| RNA-seq based | Stringtie (Illumina) | 26,700 | 705,699 |
|  | Stringtie (Iso-seq) | 5,251 | 50,443 |
|  | PASA | 176,040 | 2,194,858 |
| EVM Integration | | 28,905 | 255,740 |
| PASA Update | | 29,278 | 585,487 |

**Supplementary Table 13** Whole-genome gene annotation for *S. scherzeri* using different prediction approaches

| **Gene set** | | **Gene Number** | **Exon Number** |
| --- | --- | --- | --- |
| *Ab initio* based | Braker2 (Illumina) | 39,599 | 396,977 |
|  | Braker2 (PacBio Iso-seq) | 42,435 | 271,365 |
| Homolog based | *Danio rerio* | 30,977 | 291,720 |
|  | *Oryzias latipes* | 41,571 | 357,174 |
|  | *Takifugu rubripes* | 25,743 | 253,974 |
|  | *Gasterosteus aculeatus* | 31,746 | 281,875 |
|  | *Tetraodon nigroviridi* | 21,220 | 231,206 |
| RNA-seq based | Stringtie (Illumina) | 23,844 | 741,156 |
|  | Stringtie (Iso-seq) | 11,771 | 94,293 |
|  | PASA | 135,467 | 1,718,623 |
| EVM Integration | | 29,497 | 256,599 |
| PASA Update | | 29,543 | 604,121 |

**Supplementary Table 14** BUSCO evaluation of predicted gene models of *S. chuatsi* and *S. scherzeri*

|  | *S. chuatsi* | *S. scherzeri* |
| --- | --- | --- |
| Complete BUSCOs | 3353 (92.1%) | 3337 (91.7%) |
| Complete and single-copy BUSCOs | 2136 (58.7%) | 2158 (59.3%) |
| Complete and duplicated BUSCOs | 1217 (33.4%) | 1179 (32.4%) |
| Fragmented BUSCOs | 44 | 73 |
| Missing BUSCOs | 243 | 230 |
| Total BUSCO groups searched | 3640 | 3640 |

**Supplementary Table 15** Summary of gene functional annotations

|  |  | ***S. chuatsi*** | ***S. scherzeri*** |
| --- | --- | --- | --- |
| **Total** | | 29,278 | 29,543 |
| **Annotated** | InterPro | 25,507 (87.12%) | 25,899 (87.67%) |
|  | NCBI NR | 23,064 (78.78%) | 23,444 (79.36%) |
|  | KEGG | 16,119 (55.05%) | 16,544 (56.00%) |
|  | Swiss-Prot | 20,051 (68.48%) | 20,501(69.39%) |
|  | eggNOG | 20,796 (71.03%) | 21,192 (71.73%) |
|  | Total | 26,623 (90.93%) | 27,024 (91.47%) |
| **Unannotated** | | 2,655 (9.07%) | 2,519 (8.53%) |

**Supplementary Table 16** Summary of *S. chuatsi* and *S. scherzeri* individuals for resequencing

| **Sample name** | **species** | **Sampling site** | **Tissue of samples** | **Total bases (Gb)** | **Sequencing depth (**$\boldsymbol{\times}$**)** |
| --- | --- | --- | --- | --- | --- |
| SinChu1 | *S. chuatsi* | Hunan | Dorsal Fin | 8.70 | 12.15 |
| SinChu2 | *S. chuatsi* | Hunan | Dorsal Fin | 9.53 | 13.31 |
| SinChu3 | *S. chuatsi* | Hunan | Dorsal Fin | 8.84 | 12.34 |
| SinChu4 | *S. chuatsi* | Hunan | Dorsal Fin | 8.19 | 11.43 |
| SinChu5 | *S. chuatsi* | Hunan | Dorsal Fin | 8.87 | 12.38 |
| SinChu6 | *S. chuatsi* | Hunan | Dorsal Fin | 8.59 | 12.00 |
| SinSch1 | *S. scherzeri* | Jilin | Dorsal Fin | 8.14 | 11.37 |
| SinSch2 | *S. scherzeri* | Jilin | Dorsal Fin | 8.69 | 12.13 |
| SinSch3 | *S. scherzeri* | Jilin | Dorsal Fin | 8.36 | 11.67 |
| SinSch4 | *S. scherzeri* | Jilin | Muscle | 9.89 | 13.80 |
| SinSch5 | *S. scherzeri* | Jilin | Muscle | 10.04 | 14.02 |
| SinSch6 | *S. scherzeri* | Jilin | Muscle | 9.54 | 13.32 |
| SinSch7 | *S. scherzeri* | Guangdong | Dorsal Fin | 9.04 | 12.62 |
| SinSch8 | *S. scherzeri* | Guangdong | Dorsal Fin | 9.57 | 13.36 |
| SinSch9 | *S. scherzeri* | Guangdong | Dorsal Fin | 9.45 | 13.19 |
| SinSch10 | *S. scherzeri* | Guangdong | Dorsal Fin | 9.33 | 13.02 |
| SinSch11 | *S. scherzeri* | Guangdong | Dorsal Fin | 8.57 | 11.96 |
| SinSch12 | *S. scherzeri* | Guangdong | Dorsal Fin | 8.57 | 11.96 |

**Supplementary Table 17** Positively selected genes (PSGs) in *S. chuatsi*

| **No.** | **Gene ID** | **Gene Name** | **Abbreviation** | **Omega** | ***P*-value** |
| --- | --- | --- | --- | --- | --- |
| 1 | SinChu_g028183 | *phosphofructokinase, platelet* | *pfkp* | 12.777 | 6.83E-12 |
| 2 | SinChu_g012873 | *Rap guanine nucleotide exchange factor (GEF) 3* | *rapgef3* | 1.543 | 9.17E-06 |
| 3 | SinChu_g008662 | *arrestin domain containing 3a* | *arrdc3a* | 1 | 1.45E-05 |
| 4 | SinChu_g005846 | *sphingomyelin phosphodiesterase 3* | *smpd3* | 1.963 | 1.92E-05 |
| 5 | SinChu_g005932 | *membrane-bound transcription factor peptidase, site 1* | *mbtps1* | 2.892 | 1.27E-04 |
| 6 | SinChu_g006396 | *tropomyosin 1* | *tpm1* | 37.621 | 1.53E-04 |
| 7 | SinChu_g014068 | *eukaryotic translation initiation factor 2-alpha kinase 3* | *eif2ak3* | 1.542 | 3.94E-04 |
| 8 | SinChu_g022725 | *ubiquitin carboxyl-terminal esterase L3* | *uchl3* | 1.603 | 2.71E-03 |
| 9 | SinChu_g019912 | *activation-induced cytidine deaminase* | *aicda* | 5.854 | 2.96E-03 |
| 10 | SinChu_g019441 | *chymotrypsinogen B* | *ctrb* | 1.632 | 5.49E-03 |
| 11 | SinChu_g022935 | *crystallin, gamma M3* | *crygm3* | 5.738 | 7.58E-03 |
| 12 | SinChu_g001073 | *titin-cap* | *tcap* | 5.131 | 7.83E-03 |
| 13 | SinChu_g016175 | *notch receptor 1* | *notch1* | 2.341 | 8.93E-03 |
| 14 | SinChu_g004528 | *peroxiredoxin like 2B* | *prxl2b* | 1.017 | 1.48E-02 |
| 15 | SinChu_g017335 | *solute carrier family 25 member 3a* | slc25a3a | 2.022 | 4.56E-02 |

**Supplementary Table 18** Positively selected genes (PSGs) in *S. scherzeri*

| **No** | **Gene ID** | **Gene Name** | **Abbreviation** | **Omega** | ***P*-value** |
| --- | --- | --- | --- | --- | --- |
| 1 | SinSch_g018173 | *Kin17 DNA and RNA binding protein* | *kin* | 16.209 | 2.032E-14 |
| 2 | SinSch_g027281 | *prolyl 4-hydroxylase, alpha polypeptide I a* | *p4ha1a* | 14.815 | 1.32E-13 |
| 3 | SinSch_g004881 | *calcium channel, voltage-dependent, L type, alpha 1D subunit, a* | *cacna1da* | 3.126 | 1.63E-06 |
| 4 | SinSch_g015059 | *EYA transcriptional coactivator and phosphatase 4* | *eya4* | 2.176 | 4.79E-06 |
| 5 | SinSch_g004606 | *mitogen-activated protein kinase 14b* | *mapk14b* | 10.548 | 1.79E-05 |
| 6 | SinSch_g021541 | *ribosomal protein S8b* | *rps8b* | 5.625 | 2.52E-05 |
| 7 | SinSch_g021470 | *integrin, beta 1a* | *itgb1a* | 1.786 | 2.57E-05 |
| 8 | SinSch_g004447 | *STIM activating enhance* | *stimate* | 6.274 | 4.06E-05 |
| 9 | SinSch_g005151 | *cadherin 22* | *cdh22* | 2.009 | 1.11E-04 |
| 10 | SinSch_g016577 | *syntaxin binding protein 1a* | *stxbp1a* | 3.702 | 8.13E-04 |
| 11 | SinSch_g004114 | *cell division cycle 42* | *cdc42* | 1.487 | 9.01E-04 |
| 12 | SinSch_g015537 | *5'-3' exoribonuclease 2* | *xrn2* | 1.457 | 1.40E-03 |
| 13 | SinSch_g002350 | *DENN domain containing 10* | *dennd10* | 3.309 | 3.15E-03 |
| 14 | SinSch_g007414 | *recombination signal binding protein for immunoglobulin kappa J region b* | *rbpjb* | 10.465 | 3.89E-03 |
| 15 | SinSch_g010804 | *chloride channel, nucleotide-sensitive, 1A* | *clns1a* | 1.226 | 4.74E-03 |
| 16 | SinSch_g001009 | *solute carrier family 25 member 17* | *slc25a17* | 3.178 | 5.36E-03 |
| 17 | SinSch_g021912 | *ring finger and CCCH-type domains 1b* | *rc3h1b* | 1.265 | 5.39E-03 |
| 18 | SinSch_g007217 | *protein phosphatase 1, regulatory (inhibitor) subunit 14Bb* | *ppp1r14bb* | 8.416 | 5.82E-03 |
| 19 | SinSch_g003579 | *N-acetylneuraminic acid synthase a* | *nansa* | 7.354 | 6.93E-03 |
| 20 | SinSch_g027795 | *elongin B* | *elob* | 26.514 | 1.03E-02 |
| 21 | SinSch_g010515 | *proteasome 20S subunit beta 4* | *psmb4* | 1.014 | 1.17E-02 |
| 22 | SinSch_g017936 | *ubiquitin-conjugating enzyme E2Nb* | *ube2nb* | 13.274 | 1.51E-02 |
| 23 | SinSch_g001810 | *protein phosphatase, Mg2+/Mn2+ dependent, 1Ba* | *ppm1ba* | 1.737 | 1.66E-02 |
| 24 | SinSch_g008885 | *bri3 binding protein* | *bri3bp* | 4.148 | 2.38E-02 |
| 25 | SinSch_g014370 | *attractin* | *atrn* | 2.439 | 4.54E-02 |

**Supplementary Table 19** Summary of sequencing and peak calling of CUT&Tag

|  | **Species** | **Mapped pair-reads** | **Mapping rate** | **PCR repeat rate** | **Uniquely mapped reads** | **Number of peaks** | **Average length of peaks (bp)** | **FRiPs** |
| --- | --- | --- | --- | --- | --- | --- | --- | --- |
| H3K27ac | *S. chuatsi* | 22,383,313 | 97.90% | 86.62% | 2,762,865 | 17,015 | 789.66 | 66.05% |
|  | *S. scherzeri* | 25,938,076 | 96.18% | 80.85% | 3,655,502 | 15,150 | 742.97 | 69.99% |
| H3K4me3 | *S. chuatsi* | 52,063,564 | 95.14% | 73.09% | 13,173,930 | 16,955 | 1061.62 | 77.32% |
|  | *S. scherzeri* | 25,585,943 | 93.21% | 39.33% | 13,785,016 | 16,239 | 1014.29 | 72.05% |
| IgG (Control) | *S. chuatsi* | 165,620 | 26.77% | 96.07% | 2,627 | 2 | 329.50 | 2.00% |
|  | *S. scherzeri* | 451,411 | 55.46% | 95.91% | 7,697 | 4 | 322.00 | 0.22% |

**Supplementary Table 20** Summary of sequencing and peak calling of ATAC-Seq

| **Sample** | **Species** | **Mapped pair-reads** | **Mapping rate** | **PCR repeat rate** | **Uniquely mapped reads** | **Number of peaks** | **Average length of peaks (bp)** | **FRiPs** |
| --- | --- | --- | --- | --- | --- | --- | --- | --- |
| Chu1 | *S. chuatsi* | 91,891,999 | 97.29% | 25.10% | 61,507,958 | 140,942 | 371.16 | 22.13% |
| Chu2 | *S. chuatsi* | 75,938,882 | 97.35% | 22.29% | 52,783,582 | 133,094 | 362.69 | 21.14% |
| Sch1 | *S. scherzeri* | 67,435,897 | 93.63% | 18.70% | 40,670,224 | 172,513 | 267.22 | 18.72% |
| Sch2 | *S. scherzeri* | 62,998,480 | 94.40% | 15.77% | 39,595,802 | 161,158 | 306.56 | 20.62% |

**Supplementary Table 21** Summary of *cis*-regulatory regions identified in the genomes of *S. chuatsi* and *S. scherzeri*

|  | **Distal enhancers** | **Promotors** | | | **Non-redundant *cis*-regulatory regions** |
| --- | --- | --- | --- | --- | --- |
|  | **H3K27ac** | **H3K4me3** | **H3K27ac** | **Total promotors** |  |
| *S. chuatsi* | 4,965 | 16,953 | 12,048 | 17,623 | 22,588 |
| *S. scherzeri* | 3,132 | 16,239 | 11,164 | 16,726 | 19,858 |

**Supplementary Table 22** Differentially expressed immune genes associated with *cis*-regulatory regions affected by SVs

| **Gene ID** | **Abbreviation** | **Gene name** | **Ortholog in Human** | **log2FC**  **(H3K27ac**  **Peak Intensity)** | **FPKM (Spleen)** | |
| --- | --- | --- | --- | --- | --- | --- |
|  |  |  |  |  | ***S. chuatsi*** | ***S. scherzeri*** |
| SinChu_g011534 | *prss16* | *serine protease 16* | *PRSS16* | -1.93689 | 27.62 | 71.85 |
| SinChu_g015362 | *lama4* | *laminin, alpha 4* | *LAMA4* | -1.15982 | 2.2 | 30.09 |
| SinChu_g020408 | *cd22* | *cd22 molecule* | *CD22* | -1.58585 | 0.63 | 1.66 |
| SinChu_g027583 | *tecpr1b* | *tectonin beta-propeller repeat containing 1b* | *TECPR1* | -9.46312 | 1.27 | 3.86 |

**Supplementary Table 23** Characterization of broad H3K4me3 peaks

|  | **Number of broad peaks** | **H3K27ac peaks** | |
| --- | --- | --- | --- |
|  |  | **Overlap** | **Percentage** |
| *S. chuatsi* | 491 | 459 | 93.50% |
| *S. scherzeri* | 481 | 444 | 92.30% |

**Supplementary Table 24** Summary of top 20 enriched KEGG pathways of genes associated with differential broad H3K4me3 peaks

|  | **Pathway ID** | **Pathway** | **Candidate genes with pathway annotation** | **All genes with pathway annotation** | ***P*-value** | **Q-value** |
| --- | --- | --- | --- | --- | --- | --- |
| 1 | ko04072 | Phospholipase D signaling pathway | 10 (12.5%) | 144 (2.29%) | 0.000012 | 0.002434 |
| 2 | ko05206 | MicroRNAs in cancer | 8 (10%) | 134 (2.13%) | 0.000275 | 0.027607 |
| 3 | ko05200 | Pathways in cancer | 15 (18.75%) | 454 (7.22%) | 0.000501 | 0.03359 |
| 4 | ko04910 | Insulin signaling pathway | 7 (8.75%) | 139 (2.21%) | 0.001833 | 0.0729 |
| 5 | ko04151 | PI3K-Akt signaling pathway | 11 (13.75%) | 318 (5.06%) | 0.002123 | 0.0729 |
| 6 | ko05231 | Choline metabolism in cancer | 6 (7.5%) | 106 (1.69%) | 0.002176 | 0.0729 |
| 7 | ko05222 | Small cell lung cancer | 5 (6.25%) | 76 (1.21%) | 0.002678 | 0.076894 |
| 8 | ko05221 | Acute myeloid leukemia | 4 (5%) | 50 (0.79%) | 0.003585 | 0.079814 |
| 9 | ko04218 | Cellular senescence | 7 (8.75%) | 157 (2.5%) | 0.003645 | 0.079814 |
| 10 | ko05202 | Transcriptional misregulation in cancers | 7 (8.75%) | 165 (2.62%) | 0.004791 | 0.079814 |
| 11 | ko04211 | Longevity regulating pathway - mammal | 5 (6.25%) | 87 (1.38%) | 0.004806 | 0.079814 |
| 12 | ko04015 | Rap1 signaling pathway | 8 (10%) | 210 (3.34%) | 0.00497 | 0.079814 |
| 13 | ko05310 | Asthma | 3 (3.75%) | 28 (0.45%) | 0.005162 | 0.079814 |
| 14 | ko04213 | Longevity regulating pathway - multiple species | 4 (5%) | 60 (0.95%) | 0.006899 | 0.099043 |
| 15 | ko05230 | Central carbon metabolism in cancer | 4 (5%) | 62 (0.99%) | 0.007742 | 0.101914 |
| 16 | ko05169 | Epstein-Barr virus infection | 7 (8.75%) | 182 (2.89%) | 0.008113 | 0.101914 |
| 17 | ko05163 | Human cytomegalovirus infection | 7 (8.75%) | 189 (3%) | 0.009887 | 0.116899 |
| 18 | ko05162 | Measles | 5 (6.25%) | 114 (1.81%) | 0.014648 | 0.163572 |
| 19 | ko04630 | Jak-STAT signaling pathway | 5 (6.25%) | 120 (1.91%) | 0.017933 | 0.186168 |
| 20 | ko04152 | AMPK signaling pathway | 5 (6.25%) | 121 (1.92%) | 0.018524 | 0.186168 |

**Supplementary Table 25** Differential broad H3K4me3 peaks associated genes that are enriched in pathways in cancer (ko05200)

| **Gene ID** | **Abbreviation** | **Gene Name** | **Ortholog in Human** | **log2FC**  **(Broad H3K4me3 Intensity)** | **log2FC**  **(Gene Expression Level)** |
| --- | --- | --- | --- | --- | --- |
| SinChu_g011111 | *egln2* | *egl-9 family hypoxia-inducible factor 2* | *EGLN2* | 1.17304 | 2.23113776 |
| SinChu_g008717 | *f2r* | *coagulation factor II (thrombin) receptor* | *F2R* | 2.08824 | 3.19522938 |
| SinChu_g023426 | *kita* | *KIT proto-oncogene, receptor tyrosine kinase a* | *KIT* | 3.00812 | 4.68452923 |
| SinChu_g001477 | *ncoa4* | *nuclear receptor coactivator 4* | *NCOA4* | 1.19068 | 2.291397069 |
| SinChu_g017824 | *ccnd2a* | *cyclin D2, a* | *CCND2* | 1.90242 | 1.93104124 |
| SinChu_g022510 | *cyc1* | *cytochrome c-1* | *CYC1* | -2.17504 | -6.141871664 |
| SinChu_g010322 | *e2f3* | *E2F transcription factor 3* | *E2F3* | -1.47952 | -2.294534034 |
| SinChu_g016271 | *pik3r1* | *phosphoinositide-3-kinase, regulatory subunit 1 (alpha)* | *PIK3R1* | -1.92386 | -1.4228252 |
